# A frequency-tagged fMRI platform for accessing spatiotemporally resolved cortical computations

**DOI:** 10.1101/2024.12.19.629518

**Authors:** Reebal W. Rafeh, Geoffrey N. Ngo, Lyle E. Muller, Ali R. Khan, Ravi S. Menon, Taylor W. Schmitz, Marieke Mur

**Affiliations:** Department of Physiology and Pharmacology, Schulich School of Medicine & Dentistry, Western University, London ON, Canada; Department of Mathematics, Faculty of Science, Western University, London ON, Canada; Fields Lab for Network Computation, Fields Institute, Toronto ON, Canada; Department of Neuroscience, The University of Texas at Dallas, Richardson TX, USA; Department of Medical Biophysics, Schulich School of Medicine & Dentistry, Western University, London ON, Canada; Centre for Functional and Metabolic Mapping, Robarts Research Institute, Western University, London ON, Canada; Department of Psychology, Faculty of Social Science, Western University, London ON, Canada; Department of Computer Science, Faculty of Science, Western University, London ON, Canada

## Abstract

Understanding how the human cortex resolves competition between concurrent sensory inputs requires access to stimulus-specific responses, their temporal coupling, and nonlinear interactions at fine spatial scales. Conventional non-invasive neuroimaging cannot resolve these signals. We introduce multiplexed frequency-tagged fMRI (ft-fMRI) to resolve responses to simultaneous visual stimuli within individual cortical vertices. Across 3T and 7T, we identify fundamental, multiplexed, and intermodulation BOLD responses. Population receptive field mapping localizes multiplexed and intermodulation signals to visual-field locations where stimulus representations overlap, while response phase distinguishes positive and negative coupling to each stimulus drive. These signatures replicate across sessions, field strengths, and frequency pairings. To investigate their computational origins, we develop a population receptive field model incorporating divisive normalization between spatially intermingled neural subpopulations. The model accurately predicts vertex-wise frequency-response patterns, including intermodulation signals and their dependence on receptive field overlap. These findings establish ft-fMRI as a means to directly test cortical computations underlying sensory competition in the human visual cortex.

## Introduction

Our everyday sensory environments contain multiple simultaneously active inputs. In the sensory cortex, these multiple inputs are represented within spatially intermingled neuronal populations, where they compete for access to downstream processing. Divisive normalization is a well-established cortical computation for resolving such competition, in which the response of a neuronal population to its preferred input is scaled by the activity of other concurrently active populations ^1,2^. This scaling yields an intrinsic mutually suppressive force which prevents sensory overload from interfering with coherent perception, cognition and action. Normalization is thought to represent a canonical computation because it is expressed across the entire cortex ^3–7^ and is conserved across species ^1,8^. Despite its fundamental importance to supporting normal brain function ^9–11^, experimental examination of normalization has historically required invasive electrophysiological recording, where access to the strength and temporal coupling of cortical responses to competing stimulus drives, and the fine-grained maps in which these cortical responses are organized, are jointly resolvable. To date, no non-invasive neuroimaging modality can currently provide all of these dimensions simultaneously.

Frequency-tagging provides a direct strategy for overcoming this obstacle. By modulating concurrent stimuli at distinct temporal frequencies, frequency-tagging assigns each input a separable temporal identity that can be recovered from the resulting neural response. This enables multiple stimulus-specific responses to be tracked simultaneously within the same measurement, while preserving information about both the magnitude and phase of their coupling to the driving inputs. Response magnitude reflects the strength of stimulus coupling, whereas response phase can distinguish populations whose activity varies with versus against the stimulus drive.

These properties have made frequency-tagging a powerful approach in EEG, MEG, and intracranial electrophysiology for studying perception and selective attention ^12,13^. Moreover, when simultaneously driven populations interact nonlinearly, frequency-tagging can reveal additional intermodulation components that are absent from the stimulus inputs themselves, providing a direct spectral signature of interactions between concurrent representations that may contribute to cortical computations ^13–17^.

However, although frequency-tagged M/EEG can resolve simultaneous stimulus-specific responses and their interactions with high temporal precision, its spatial resolution is coarse, limiting the localization of signals arising from closely intermingled cortical populations with different stimulus preferences. This spatial mixing also complicates interpretation of response phase, because signals from differently tuned populations with opposing phase relationships may be combined at the sensor level ^18–20^, obscuring whether each population varies with or against its corresponding stimulus drive.

Functional magnetic resonance imaging (fMRI) overcomes the limitations of M/EEG by providing access to a spatial scale capable of resolving cortical maps of distinct and overlapping stimulus preferences ^21–23^. However, conventional fMRI does not preserve the identity of multiple simultaneously presented stimuli. Competition is typically inferred from differences in response magnitude across separate stimulus conditions rather than resolved directly within the same measurement ^24,25^. Here, we introduce a multiplexed frequency-tagged fMRI (ft-fMRI) framework that combines the spatial specificity of fMRI with the temporal separation afforded by frequency-tagging, enabling concurrent stimulus-specific responses to be dissociated within individual cortical vertices while preserving information about both the magnitude and phase of their coupling to the corresponding stimulus drive (Fig. 1).

**Figure 1.**
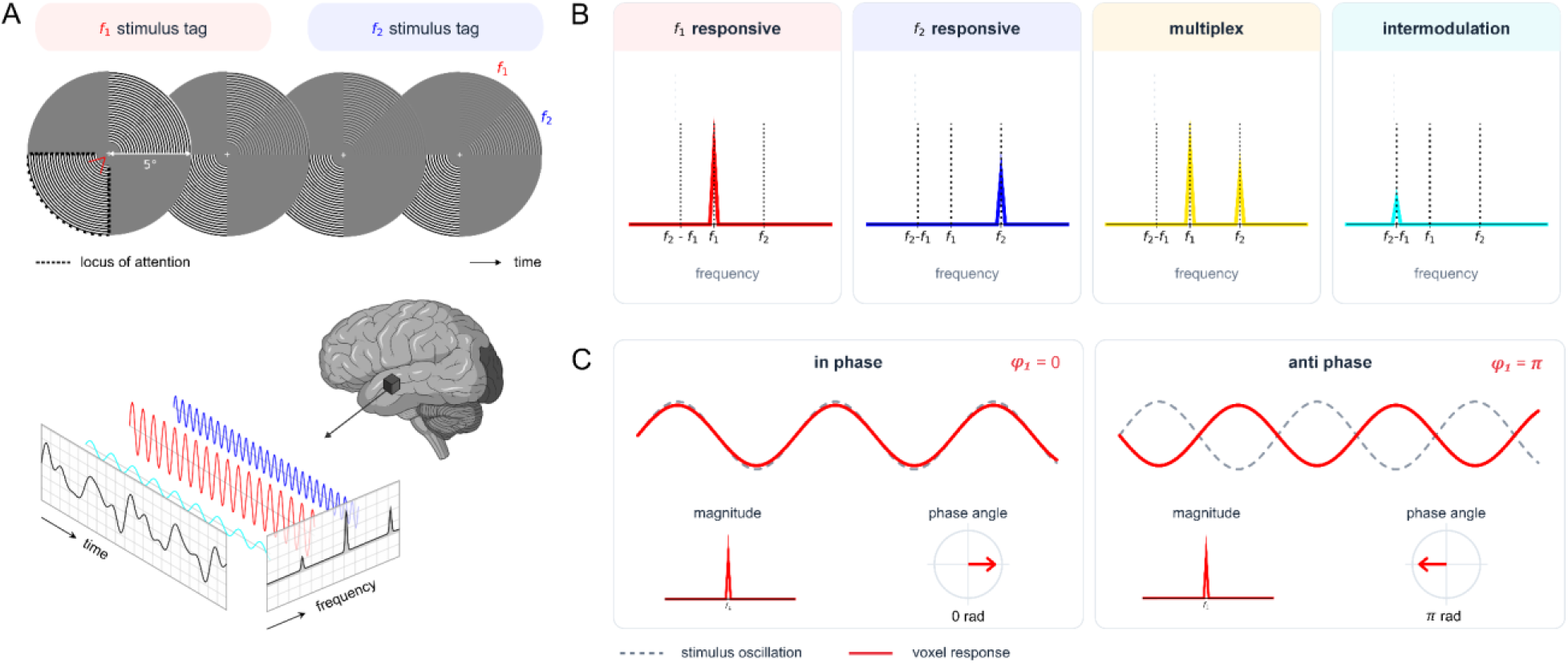
A frequency-tagged fMRI platform for spatiotemporally resolved access to cortical computations. (A) ft-fMRI stimulus configuration and schematic of frequency-tagged BOLD responses. Participants covertly attended a static radial wedge stimulus in a lower visual field quadrant, while two wedges in the opposing quadrant oscillated at distinct frequencies (*f₁* = 0.125 Hz; *f₂* = 0.2 Hz). Frequency-tagged BOLD responses are measured in individual voxels and projected onto cortical surface vertices for analysis. (B) Schematic illustrating the response components that can be resolved from individual vertex time series. Spatially dissociable neuronal populations encode the fundamental frequencies defined by the stimulus oscillations (*f₁*, red; *f₂*, blue). Vertices at the spatial overlap of these populations exhibit responses at both frequencies (multiplex, yellow). Nonlinear interactions between the fundamental frequencies give rise to an intermodulation component at the difference frequency (*f₂ − f₁*, cyan). (C) Schematic illustrating how responses at a particular frequency also provide information about temporal coupling with the stimulus. Responses may be in-phase with the stimulus oscillations, with BOLD activity increasing with stimulus intensity, or anti-phase, with BOLD activity decreasing with stimulus intensity.

Using dual-frequency visual stimulation at 3T and 7T, we identify cortical populations synchronized selectively to individual stimulus frequencies, populations multiplexing responses to both frequencies, and intermodulation components that are absent from the stimulus inputs and reflect nonlinear interactions between concurrent responses. We further show that the response phase provides complementary information about the direction of stimulus coupling. These frequency- and phase-resolved response maps are reproducible within individuals across sessions, field strengths, and frequency pairings.

We then apply population receptive field mapping to precisely localize these frequency- and phase-resolved signals according to the spatial organization of their stimulus-specific preferences. By combining these ingredients, we directly access the spatiotemporally resolved signals supporting normalization. Our results suggest that enhancement driven by one stimulus is modulated by suppression associated with the competing stimulus, giving rise to intermodulation signals. We then developed a novel population repetitive field model incorporating sub-voxel suppressive interactions accessible only with ft-fMRI. This model reproduced the intermodulation responses observed empirically from our frequency- and phase-resolved maps, whereas a model lacking these interactions did not. Together, these findings establish ft-fMRI as a framework for non-invasively resolving concurrent cortical representations, their stimulus-linked enhancement and suppression, and the nonlinear interactions through which divisive normalization may be implemented. Looking ahead, ft-fMRI provides the core ingredients needed for non-invasively accessing the mesoscale neural computations supporting cognition in naturalistic sensory environments.

## Results

### Frequency tagging reveals fundamental and multiplexed BOLD responses in human visual cortex

We estimated stimulus-evoked BOLD responses using a multiplexed frequency-tagging fMRI design. Participants (n = 11 at 3T; n = 6 at 7T) performed a covert attention task in which they detected brief color changes in a static grating presented in the lower left or right visual hemifield while maintaining central fixation (Fig. 1a; Supplementary Video 1). Concurrently, two unattended gratings were presented in the contralateral upper visual hemifield. The spatial configuration of attended and unattended stimuli was counterbalanced by hemifield across participants (see example participants Fig. 2). Behavioral performance and eye-tracking confirmed task engagement and stable fixation throughout scanning (Supplementary Fig. 1).

**Figure 2.**
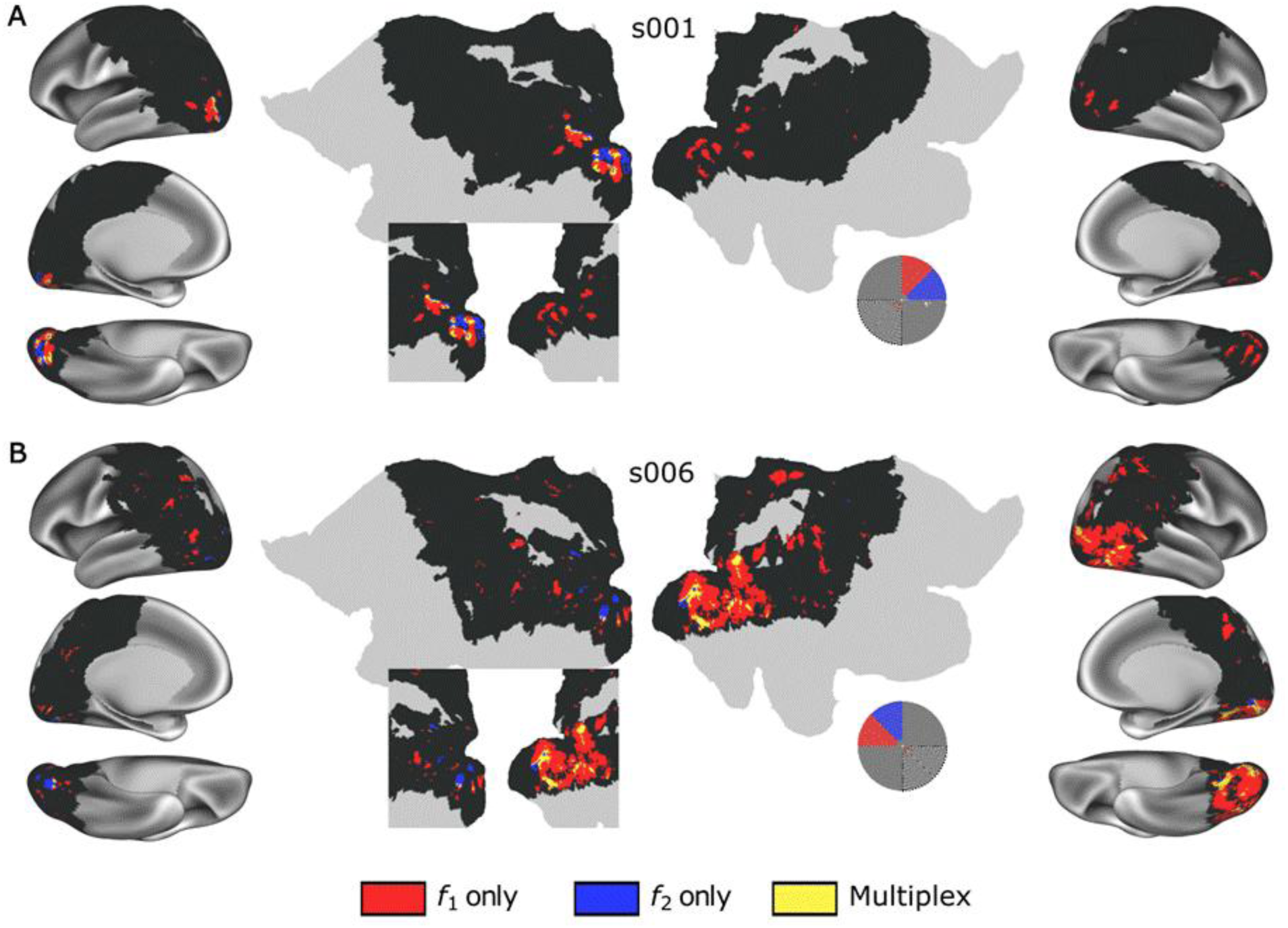
Dual frequency paradigm enables encoding of *f₁*, *f₂*, and multiplexed combinations in the visual cortex. Frequency-encoded maps were estimated in individual participants using a Monte Carlo subsampling procedure applied to the frequency-tagging condition (*n* = 400 subsamples). Vertices were considered significant at *P* < 0.05 and retained if identified in ≥ 80% of subsamples. Red and blue indicate vertices encoding *f₁* and *f₂*, respectively, and yellow indicates vertices exhibiting multiplexed responses to both frequencies. Insets show the stimulated visual field locations for *f₁* (red) and *f₂* (blue), which was counterbalanced across participants to confirm hemispheric activation was contralateral to visual stimulation. (A) Whole-brain and zoomed-in maps for a representative participant (s001) in the 3T experiment with stimulation in the upper-right visual field. (B) Same as in panel A, for a different participant (s006) with stimulation in the upper-left visual field.

To drive frequency-tagged BOLD responses at distinct frequencies, the luminance of unattended gratings was continuously modulated between visible and invisible states at two temporal frequencies (*f_1_* and *f_2_*; 0.125 and 0.2 Hz in Experiment 1; Fig. 1b, Supplementary Video 1). To capture fast neuronally driven BOLD dynamics, fMRI data were acquired at TR = 300 ms, consistent with prior work on rapid frequency-tagged BOLD responses^26–28^. Acquisition focused on a slab encompassing occipital and inferior parietal cortices.

We first identified frequency-tagged BOLD responses by fitting sine and cosine regressors for each stimulated frequency using a vertex-wise general linear model (GLM), which additionally controlled for motion and physiological noise (Methods). A Monte Carlo subsampling procedure generated robust spatial maps of frequency encoding, retaining vertices that exceeded an uncorrected threshold (P < 0.05) in at least 80% of subsampled runs. We focused first on visual cortical vertices activated in-phase with the frequency-tagged stimuli, i.e. those rising and falling with the stimulus ^29–31^ (Fig. 2, Supplementary Fig. 2), which constitute the most widely studied BOLD responses in conventional fMRI studies.

Because tagging frequencies are defined a priori by stimulus timing, responses can also be isolated directly in the frequency domain ^29,30^. To test this, we performed nuisance signal regression on the time series using a GLM and then quantified spectral power at the expected tagging frequencies relative to adjacent bins (Fourier z-score > 3)^29,30^. This spectral analysis reproduced the spatial distribution and selectivity patterns obtained with sine–cosine regression (Supplementary Fig. 3), indicating convergence between time-domain and frequency-domain inference.

Robust frequency-tagged responses were observed at both driving frequencies throughout the visual cortex (Fig. 2, Supplementary Fig. 2). As expected from the counterbalanced hemifield stimulation, responses were stronger in the visual cortex contralateral to the stimulus. Across experiments, both frequencies localized reliably to early and intermediate visual areas. Spatial maps revealed two largely distinct cortical populations preferentially encoding either *f_1_* or *f_2_*, and modulation strength tracked the counterbalanced stimulus locations between and within hemifields (e.g., subjects 001 vs. 006 in Fig. 2; Supplementary Fig. 2). Consistent with viscoelastic constraints on the BOLD response, sensitivity to the higher frequency (*f_2_*) was reduced relative to the lower frequency (*f_1_*) (P < .05; Wilcoxon signed-rank test; Supplementary Fig. 4), in line with prior reports^26,32^.

To verify that these responses were stimulus-locked rather than reflecting endogenous fluctuations or noise, participants also completed a control condition in which all stimuli were static (Supplementary Fig. 5). Frequency-tagged responses were nearly abolished in this condition (mean vertex count: 33 for *f_1_*, 3 for *f_2_*; P < .05), and any residual activity was spatially sparse and inconsistent across participants. In a subset of participants (n = 8), EEG recordings under an identical setup confirmed that the fundamental frequencies were present in occipital electrodes and exhibited physiological delays consistent with the BOLD responses (Supplementary Fig. 6), supporting a neuronal origin.

We next asked whether a single cortical vertex could simultaneously encode both stimulus frequencies, i.e., multiplexing, consistent with access to BOLD dynamics of two feature-tuned neuronal populations within the same vertex. Using the same subsampling framework, we identified individual vertices exhibiting significant frequency-tagged responses at both *f_1_* and *f_2_* (Fig. 2). These multiplexed responses were absent in the static control condition (Supplementary Fig. 5a), indicating that they arise specifically from concurrent rhythmic stimulation. Together, these results demonstrate that ft-fMRI can resolve both frequency-specific and multiplexed stimulus-driven BOLD responses within individual voxels across the human visual cortex.

We next conducted a population receptive field (pRF) mapping experiment^22^ in a subset of participants (n = 9) to estimate the visual field preferences of frequency-specific and multiplexing vertices (Fig. 3; Supplementary Fig. 7). We reasoned that vertices synchronized to a single stimulus frequency should preferentially represent the spatial location of that stimulus, whereas multiplexing vertices should represent the locations of both concurrently presented stimuli. To test this prediction, we first computed visual field coverage maps for frequency-specific and multiplexing vertices, to show what parts of the visual field these vertices respond to in the pRF mapping experiment (Fig. 3A). We observed that the pRFs of *f_1_*-encoding vertices exhibited overlap with the physical location of the *f_1_* stimulus, while *f_2_*-encoding vertices exhibited overlap with the physical location of the *f_2_* stimulus (Fig. 3B). Furthermore, pRFs of multiplexing vertices fell primarily along the diagonal boundary of the stimulus configuration, where the two stimuli are spatially adjacent (Fig. 3C). To confirm this observation, we quantified, for each vertex group, the relative overlap between its pRF coverage and the spatial location of the two frequency-tagged stimuli by computing a signed overlap index (overlap*_f1_* − overlap*_f2_*). Consistent with retinotopic organization of frequency-tagged responses, pRFs synchronized selectively to *f_1_* exhibited a bias toward the *f_1_* stimulus location (Wilcoxon signed-rank test versus 0: *P_two-tailed_*=0.027; D=1.123), while pRFs synchronized selectively to *f_2_* exhibited a bias towards the *f_2_* stimulus location (Wilcoxon signed-rank test versus 0: *P_two-tailed_*=0.0039; D=2.634). Multiplexing pRFs exhibited negligible bias towards either stimulus, indicating overlapping representation of both stimulus locations (Wilcoxon signed-rank test versus 0: *P_two-tailed_*=0.129; D=0.625). The response biases of *f_1_*, *f_2_* and multiplex vertices were also significantly different from one another (Fig 3D, Wilcoxon rank sum tests: *P_two-tailed_*<0.05 for all pairwise comparisons).

**Figure 3.**
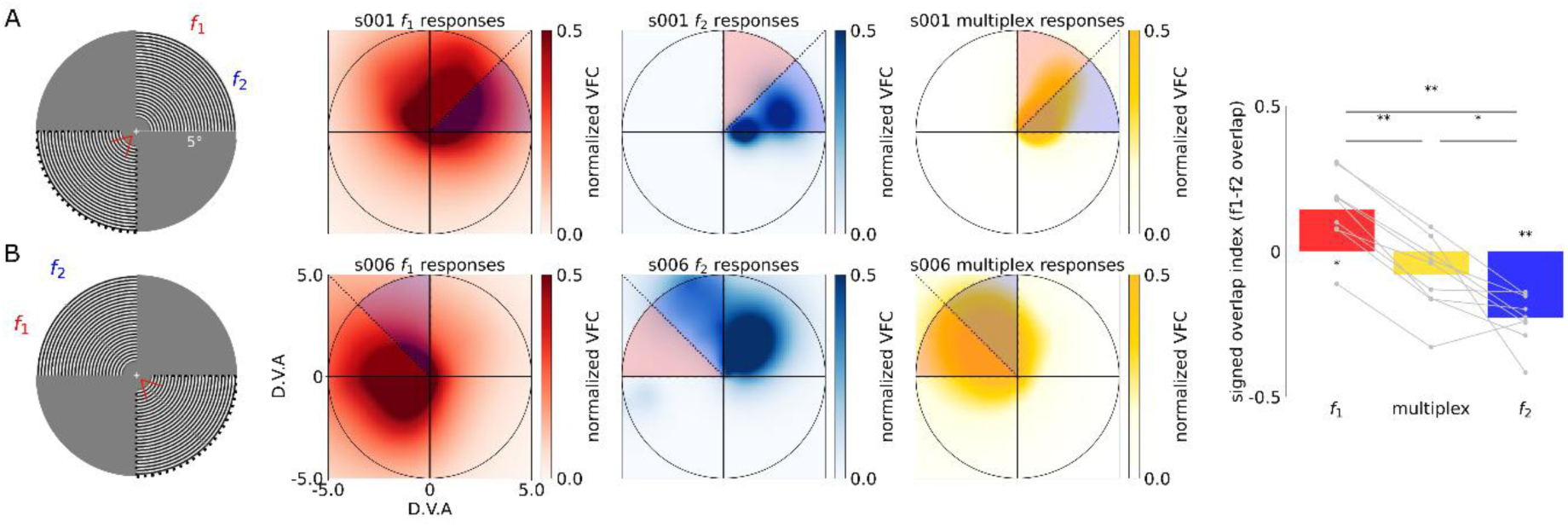
Frequency-tagged responses exhibit retinotopic preferences for the locations of their corresponding stimuli. A) Visual field coverage maps of frequency-specific (*f1*: red; *f2*: blue) and multiplexing (yellow) responses for two representative participants (rows; n=2). Red wedges indicate the position and extent of the *f1* checkerboard wedge stimulus for each participant, and blue wedges indicate the position of the *f2* wedge for each participant. Normalized visual field coverage (VFC) represents the proportion of pRFs responding to a visual field location. D.V.A is degrees visual angle. B) Visual field bias towards the location of the frequency-tagged stimuli quantified using signed overlap index. A positive value indicates greater overlap with the location of the *f1* stimulus than the *f2* stimulus. A negative value indicates greater overlap with than the location of the *f2* stimulus. A value of zero indicates equal overlap with the *f1* and *f2* stimulus locations. Statistical significance was determined using a Wilcoxon signed-rank test (\**Ptwo-tailed* < 0.05; \*\**Ptwo-tailed* < 0.01).

### Frequency-tagged fMRI reveals internally generated neural interactions in human visual cortex

In nonlinear systems such as the brain^33^, simultaneous frequency-tagged inputs can generate additional frequency components—intermodulation (IM) components—such as *f_2_* − *f_1_*, *f_1_* + *f_2_*, 2*f_1_* − *f_2_*, and 2*f_2_* − *f_1_*^14,34^. In electrophysiological studies, these components are widely interpreted as signatures of lateral and feedback processing underlying perception and attention^15,35,36^. Whether such internally generated signals are detectable with fMRI remains unexplored.

To test whether ft-fMRI is sensitive to IM, we first examined four IM components that have frequencies accessible with ft-fMRI (*f_2_* − *f_1_* = 0.075 Hz, *f_1_* + *f_2_* = 0.325 Hz, 2*f_1_* − *f_2_* = 0.05 Hz, 2*f_2_* − *f_1_* = 0.275 Hz). This was accomplished using the same GLM-based Monte Carlo subsampling procedure as before for fundamental frequencies (Fig. 2). We observed vertices exhibiting frequency-specific responses for all four of the IM components. Of these four components, the number of vertices expressing the *f_2_*-*f_1_* (0.075Hz) component most strongly differentiated the frequency-tagging from the static control condition (Fig. 4A). Furthermore, we found that these increases were localized to the hemisphere contralateral to the location of frequency-tagged stimuli (Fig. 4B; Supplementary Fig. 8,9). Altogether, these results indicate that IM is detectable and tracks with the experimental manipulation of our visual stimulation paradigm.

**Figure 4.**
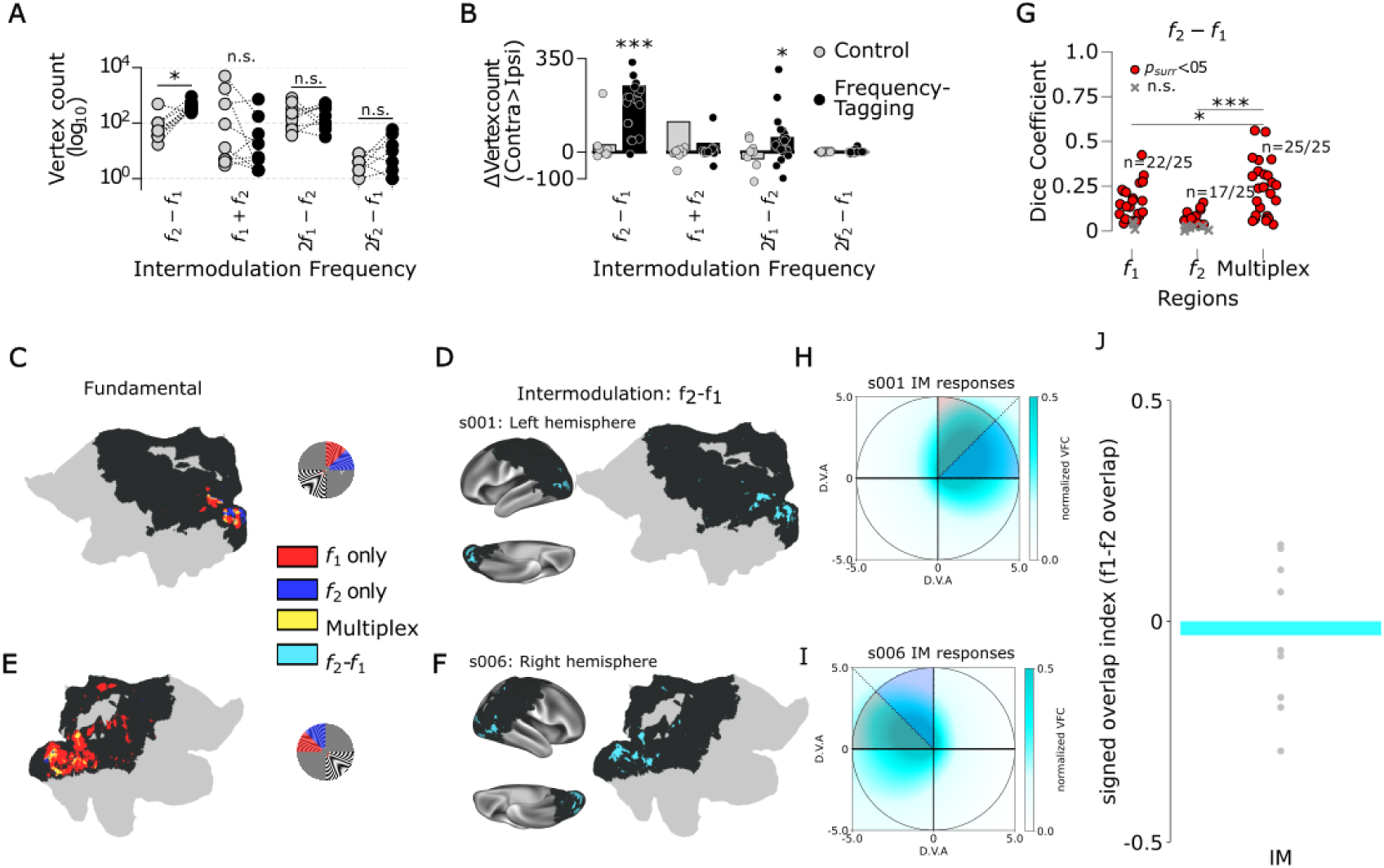
Nonlinear intermodulation responses emerge in cortical populations jointly encoding both frequency-tagged stimuli. A) Number of localized vertices at each IM frequency during the frequency-tagging (black data points) versus static control (gray data points) condition. Each data point represent the number of vertices meeting the thresholding criteria (fractional overlap > 0.8; *Punadjusted* < 0.05) for an individual participant. B) The difference in the number of IM vertices localized in the cortical hemisphere contralateral to frequency-tagged stimulus presentation and the hemisphere ipsilateral to stimulus presentation. Data points follow the same conventions as in (A) C) Localized linear fundamental frequencies on the cortical surface contralateral to the frequency-tagged stimulus presentation for a representative participant. Stimulus configuration icon depicts the spatial location of *f₁* (red wedge) and *f₂* (blue wedge) presentation. D) Localized non-linear intermodulation frequency (*f₂* - *f₁*) on the cortical surface contralateral to the frequency-tagged stimulus presentation for a representative participant. E) Data follows the same convention as in (C), except that the frequency-tagged stimulus presentation was reversed for this representative participant. F) Data follows the same convention as in (D), except that the frequency-tagged stimulus presentation was reversed for this representative participant. G) Spatial overlap between frequency-specific and multiplex, and IM responses quantified using Dice Coefficient. Statistical significance of overlap was determined using a surrogate map test. Significant differences in overlap across vertex groups were determined using a Wilcoxon signed-rank test (\**Pone-tailed* < 0.05; \*\*\**Pone-tailed* < 0.01). H) Visual field coverage maps of IM responses for representative participant shown in panels (C) and (D). Red wedges indicate the position and extent of the *f₁* checkerboard wedge stimulus for each participant, and blue wedges indicate the position of the *f₂* wedge for each participant. Normalized visual field coverage (VFC) represents the proportion of IM pRFs responding to a visual field location. D.V.A is degrees visual angle. I) Visual field coverage maps of IM responses for representative participant shown in panels (E) and (F). Data follows the same conventions as in (H). J) Visual field bias towards the location of the frequency-tagged stimuli quantified using signed overlap index. A positive value indicates greater overlap with the location of the *f₁* stimulus than the *f₂* stimulus. A negative value indicates greater overlap with than the location of the *f₂* stimulus. A value of zero indicates equal overlap with the *f₁* and *f₂* stimulus locations. Statistical significance was determined using a Wilcoxon signed-rank test (\**Ptwo-tailed* < 0.05; \*\**Ptwo-tailed* < 0.01).

We next investigated how IM responses relate spatially to the cortical populations encoding the fundamental frequencies (*f_1_*, *f_2_*, multiplex) across individuals. For each subject, we found that the locations of their IM responses track with the cortical distribution of fundamental frequency responses (Fig. 4C,D; Supplementary Fig. 7D). Consistent with this observation, intrasubject overlap analyses revealed that *f_2_* − *f_1_* maps overlapped significantly with *f_1_*-only populations in 22/25 experiments, *f_2_*-only populations in 17/25 experiments, and multiplexed populations in all 25 experiments (P < .05; surrogate map test; Fig. 5E). Although IM vertices reliably exhibited above chance overlap with *f_1_*, *f_2_*, and multiplex vertices within individuals, the magnitude of this overlap was significantly larger for multiplexing compared to fundamental frequency vertices (P < .05; Wilcoxon signed-rank test).

**Figure 5.**
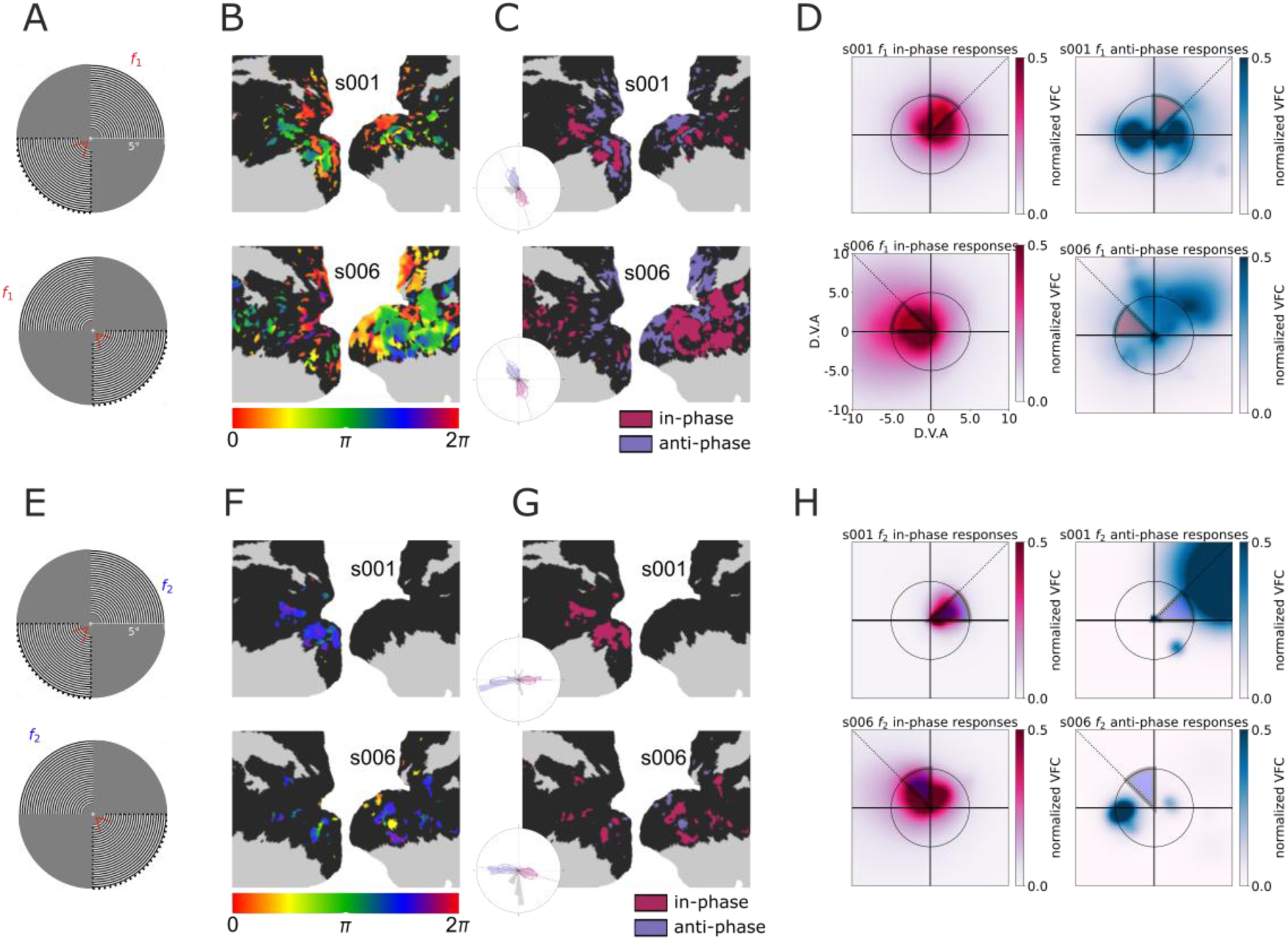
Frequency-tagged responses exhibit stimulus-linked enhancement and suppression. A) *f1* stimulus configuration for representative participants (s001 and s006; rows). Surface and polar distribution data in subsequent panels (B-D) are based on these participants. B) Phase maps for suprathreshold *f1* (fractional overlap>0.8) visual cortical vertices. Color bars denote the response phase scaled to the period of the examined frequency (1/*f1*). C) Polar plots display the response phase distribution across suprathreshold *f1* visual cortical vertices. Colored vertices consistently responded in-phase (violet-red) or anti-phase (slate-blue) with the stimulus oscillations across subsamples of the data (>0.95 of subsamples). Signed phase maps show the locations of these vertices on the cortical surface. Vertices dissociate into spatially distinct distributions. D) Visual field coverage maps of suprathreshold in-phase (violet-red) and anti-phase (slate-blue) vertices at *f1*. Outlined red wedges indicate the position and extent of the *f1* checkerboard wedge stimulus for each participant. Normalized visual field coverage (VFC) indicates the proportion of pRFs responding to a visual field location. D.V.A is degrees visual angle. The stimulus configuration for each participant is shown on the left. E) *f2* stimulus configuration for representative participants (s001 and s006; rows). Surface and polar distribution data in subsequent panels (F-H) are based on these participants. F) Phase maps for suprathreshold *f2* (fractional overlap>0.8) visual cortical vertices. Color bars denote the response phase scaled to the period of the examined frequency (1/*f2*). G) Phase distribution and signed phase maps for suprathreshold *f2* visual cortical vertices. Data organization follows the same conventions as panel C. H) Visual field coverage maps of suprathreshold in-phase (violet-red) and anti-phase (slate-blue) vertices at *f2*. Outlined blue wedges indicate the position and extent of the *f2* checkerboard wedge stimulus for each participant. Data organization follows the same conventions as in panel D.

If IM is spatially constrained by multiplexing vertices, then we would expect from our pRF analysis that the signed overlap index of IM vertices, like multiplex vertices, should be centered around zero, indicative of spatial tuning to the locations of both frequency-tagged stimuli (Fig. 4F; Supplementary Fig. 7). Consistent with this prediction, we found that IM pRFs fell along the diagonal between the two frequency-tagged stimuli (Wilcoxon signed-rank test; *P_two-tailed_*=0.652; D=0.186). Altogether, the intra-subject overlap and pRF analysis demonstrate that nonlinear IM BOLD responses are constrained by the cortical populations jointly encoding multiple stimulus frequencies.

### Frequency-tagged fMRI reveals responses in-phase and anti-phase with the stimulus drive

The spatial confinement of intermodulation responses to cortical populations jointly encoding the two frequency-tagged stimuli suggests that these responses arise from interactions between concurrently driven neural populations. To characterize these interactions further, we considered an additional dimension of the frequency-tagged response: its phase relative to the corresponding stimulus drive (Fig. 1C). Whereas response magnitude quantifies the strength of coupling to a tagged stimulus, response phase indicates the sign of that temporal coupling. So far we have focused on responses in-phase with the stimulus oscillations, which are generally interpreted as stimulus-driven activation ^21,22,29,30,37,38^. However, prior work has also shown responses synchronized anti-phase with stimulus oscillations, which have been interpreted as stimulus-driven suppression ^37–40^. Because ft-fMRI separates responses associated with different simultaneously presented frequencies, these phase relationships can be mapped independently for each stimulus within the same cortical measurement ^41^.

We estimated the response phase within our GLM approach as the angle between the sine and cosine model fits for each fundamental frequency in suprathreshold vertices (fractional overlap > 0.8). Phase maps show spatial structure that is similar across participants who viewed the same stimulus configuration (Fig. 5A,B; Supplementary Fig. 10,11). Across the visual cortex, phase estimates were concentrated predominantly around two opposing phase relationships: in-phase and anti-phase with the stimulus oscillations (Fig. 5B,D insets; Supplementary Fig. 12,13). In-phase responses increased as the corresponding stimulus drive increased, whereas anti-phase responses showed the inverse relationship, decreasing as stimulus drive increased. These response classes also exhibited distinct spatial organization (Fig. 5C,E; Supplementary Fig. 14,15). In-phase responses preferentially represented the location of the corresponding frequency-tagged stimulus, whereas anti-phase responses preferentially represented surrounding regions of visual space (Fig. 5F,G;Supplementary Fig. 16). Thus, phase maps reveal spatially organized populations that are positively versus negatively coupled to the same stimulus drive ^41^.

### Frequency-tagged BOLD responses are stable within subject, robust to MR field strength, and frequency spacing

We first asked whether fundamental (*f_1_*, *f_2_*) and intermodulation (IM; *f_2_*−*f_1_*) responses could be observed directly in the BOLD time series and maintained over repeated acquisitions within an individual. In a representative participant sampled over 24 runs, carpet plots of frequency-selective vertices revealed prominent periodic oscillations at the corresponding frequencies, confirmed by peaks in their power spectra (Fig. 6D–F). These oscillations remained clearly visible within individual three-minute runs (Fig. 6G–I), while fractional-overlap maps demonstrated consistent spatial encoding of *f_1_*, *f_2_*, and IM across the 24 runs, with the greatest sensitivity at *f_1_* (Fig. 6A–C). Both fundamental frequencies were also evident in the spectra of multiplexing vertices. Thus, the separable spectral components of ft-fMRI can be observed directly in BOLD dynamics and remain detectable across extensive repeated sampling.

**Figure 6.**
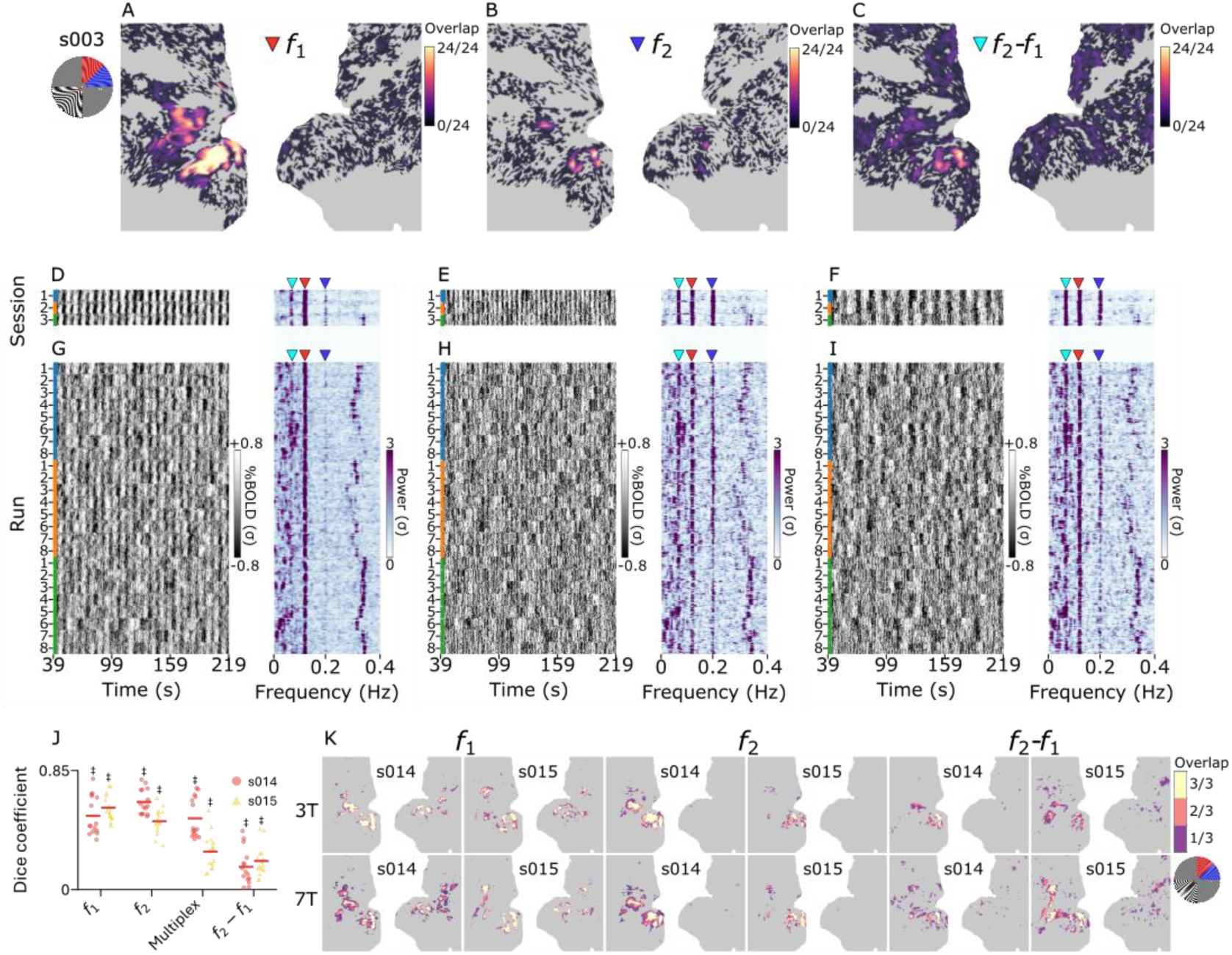
Frequency-tagged BOLD responses are stable within subject, robust to MR field strength, and frequency spacing. A) Overlap between single run localized fundamental frequency (*f₁*) responses in a representative participant. Overlap indicates the proportion of runs that vertices met the *Punadjusted* < 0.05 threshold. B) Data follows the same conventions as in panel (A), but for *f*₂. C) Data follows the same conventions as in panel (A), but for IM (*f₂* - *f₁*). D) Left: Individual session carpet plots (rows) showing the timecourse (x-axis) of frequency-tagged vertices (y-axis) occurring in at least 50% of single runs at *f₁*. Right: Power spectra corresponding to the carpet plots. Frequencies of interest indicated by colored triangles (*f₁:* red*; f₂:* blue*; f₂* - *f₁*: cyan) E) Data follows the same conventions as in panel (D), but for *f₂* vertices. F) Data follows the same convention as in (D), but for IM (*f₂* - *f₁*) vertices G) Left: Individual run carpet plots (rows) showing the timecourse (x-axis) of frequency-tagged vertices (y-axis) at *f₁*. Right: Power spectra corresponding to the carpet plots. Frequencies of interest indicated by colored triangles (*f₁:* red*; f₂:* blue*; f₂* - *f₁*: cyan) H) Data follows the same conventions as in panel (G), but for *f₂* vertices. I) Data follows the same conventions as in panel (G), but for *f₂* - *f₁* vertices. J) Overlap between frequency-specific maps (fractional overlap > 0.8) associated with different frequency pairings (n=3) and field strengths (n=2) for s014 (red data points) and s015 (purple data points) quantified using Dice’s Coefficient. (One sampled t-test; ‡P < 0.001). K) Visualization of the overlap maps across frequency pairings at each frequency of interest (*f₁:* left*; f₂:* middle*; f₂* - *f₁*: right) for the different subjects (columns) at different field strengths (3T and 7T: rows)

We next quantified the spatial reproducibility of these responses within and between individuals. Dice overlap coefficients showed significantly greater within-than between-individual spatial correspondence for *f_1_*, *f_2_*, multiplex, and IM maps at both run and session levels (Mann–Whitney U-test: *P* < 0.05; Supplementary Fig. 17). Phase maps similarly exhibited significantly greater within-than between-individual circular correlations across sessions (*P* < 0.05; Supplementary Fig. 18). Together, these results establish reproducible, individual-specific spatial organization of both frequency- and phase-resolved responses.

To test generalizability across acquisition conditions, we examined three frequency pairings in two observers scanned at both 3T and 7T, systematically varying either *f_1_* or *f_2_* in steps of 0.025 Hz (s014: *f_1_* = 0.125, 0.15, 0.175 Hz with fixed *f_2_* = 0.2 Hz; s015: *f_2_* = 0.2, 0.175, 0.15 Hz with fixed *f_1_* = 0.125 Hz) Spatial maps of *f_1_*, *f_2_*, multiplex, and IM responses remained consistent across frequency pairs and field strengths (*P* < 0.05; one-sample t-test; Fig. 6J–K). Notably, IM maps remained spatially reproducible despite changes in the underlying fundamental frequencies, indicating that their localization was not specific to a particular frequency pairing.

Together, these findings demonstrate that fundamental, multiplexed, IM, and phase-resolved BOLD responses exhibit reproducible spatial organization within individuals and remain robust across the tested acquisition conditions. Having established the reliability of these complementary response dimensions, we next asked whether they could provide access to the computations governing interactions between concurrent cortical representations.

### Leveraging frequency-tagged BOLD responses to access population-level cortical computations

Two complementary properties of frequency-tagged BOLD responses provide a potential window into cortical computation. First, intermodulation (IM) responses emerge at frequencies absent from the stimulus inputs, revealing nonlinear response components preferentially localized to cortical populations jointly encoding both stimuli. Second, response phase distinguishes populations whose activity varies with versus against the corresponding stimulus drive. These opposing phase relationships exhibit a spatially organized target–surround arrangement, with in-phase responses preferentially representing the stimulus location and anti-phase responses representing surrounding regions of visual space. Together, these observations suggest that IM may reflect nonlinear interactions between stimulus-driven enhancement and suppression within overlapping cortical representations.

Divisive normalization provides a candidate computational mechanism linking these observations. Under normalization, the response of a neuronal population to its preferred stimulus is scaled by activity associated with competing inputs ^1,2,42^. Such interactions could produce nonlinear mixing of concurrent stimulus drives, giving rise to IM components absent from the physical inputs. However, although we have shown that IM responses preferentially localize to multiplexing vertices whose receptive fields overlap both stimuli, this spatial correspondence alone does not establish how the constituent neural populations interact to generate IM. We therefore asked whether suppressive interactions between competing neuronal populations could account for the observed nonlinear response components.

To address this question, we extended established population receptive field (pRF) frameworks ^21,22^, which typically characterize the spatial tuning of each fMRI vertex using a single aggregate receptive field. Because frequency-tagging enables the contributions of distinct stimulus-selective populations to be dissociated within the same single-voxel BOLD measurement, we developed a sub-voxel pRF framework that explicitly models multiple constituent neuronal populations and their potential interactions (Supplementary Video 2; Fig. 7). This approach allowed us to distinguish the consequences of independent stimulus-driven responses from those arising through suppressive interactions between spatially overlapping populations.

**Figure 7.**
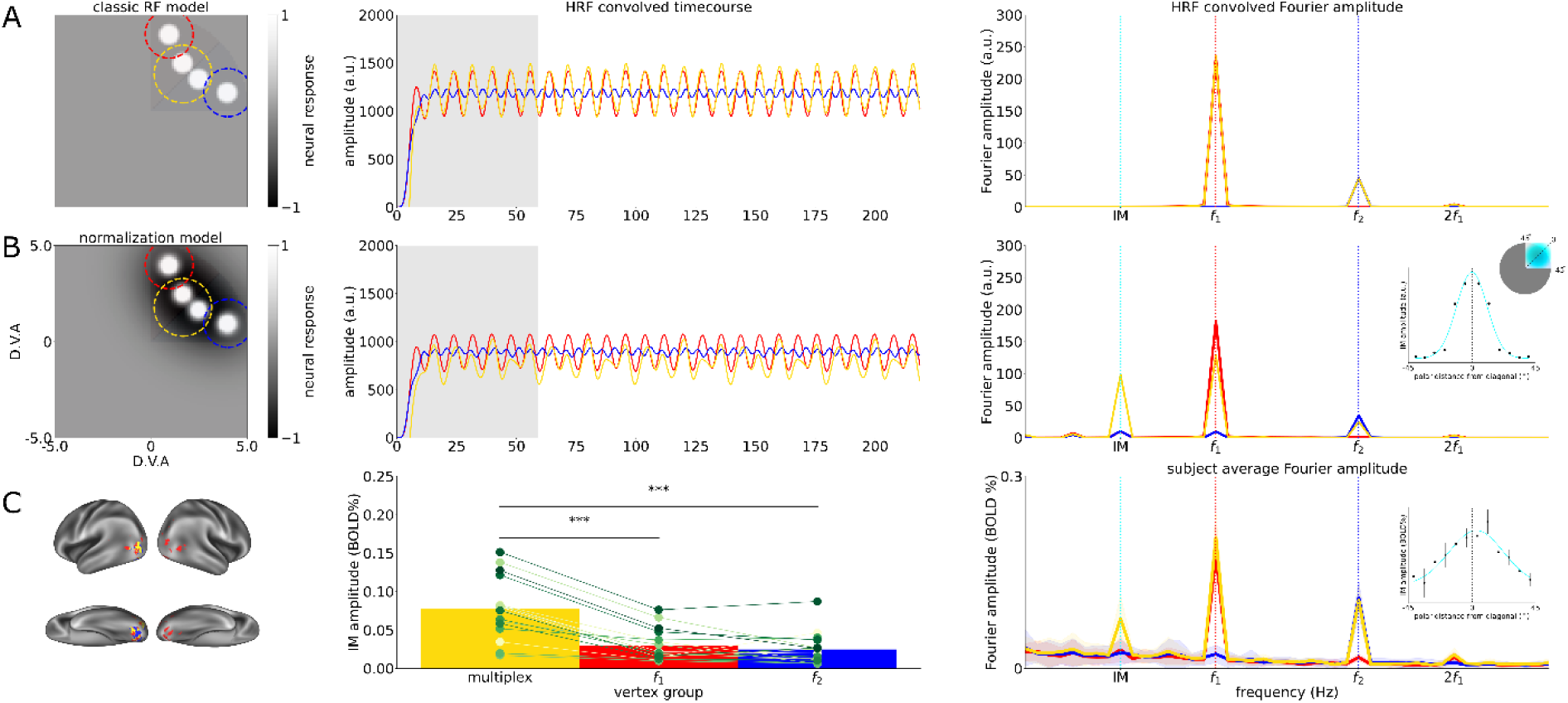
Leveraging frequency-tagged BOLD responses to access population-level cortical computations. A) **Left.** Simulated classic neural responses of differentially tuned neural subpopulations responding to the *f₁* stimulus (red wedge, dashed outline), *f₂* stimulus (blue wedge, dashed outline), or multiplexing both stimuli (yellow dashed outline). **Middle**. HRF convolved time course of subpopulation responses under our frequency-tagging design. The shaded gray area represents the time window omitted from Fourier analysis to account for hemodynamic response onset following frequency-tagged stimulation. **Right**. Fast Fourier Transform amplitude of the simulated HRF convolved time course, with dashed vertical lines indicating the IM frequency (cyan) and stimulus frequencies (*f₁*: red; *f₂*: blue) in the amplitude spectrum. B) Simulated normalization model responses under our frequency-tagging paradigm. The normalization model introduces a suppressive field to each subpopulation. Data organization follows the same conventions as in panel (a). Inset in the right panel indicates simulated changes in intermodulation amplitude as a function of polar distance from the diagonal between the frequency-tagged stimulus representations. Black data points indicate simulated IM amplitude at a given polar distance from the diagonal. C) **Left:** Frequency-tagged responses on the cortical surface of a representative participant. **Middle:** average IM BOLD amplitude in multiplexing and frequency-specific vertices across participants. Data point colors represent individual participants. **Right:** Participant average spectrum of brain responses in multiplexing and singular voxels. Shaded area represents standard deviation across sessions. Inset in the right panel indicates observed changes in intermodulation amplitude as a function of polar distance from the diagonal between the frequency-tagged stimulus representations. Black data points indicate measured IM amplitude at binned angular distances from the diagonal. Error bars indicate standard error across IM amplitude estimates within a bin. Statistical significance was determined using a Wilcoxon signed-rank test (\*\*\**Ptwo-tailed* < 0.001).

Within this framework, we compared two mechanistic accounts. In the feedforward model, stimulus-specific Gaussian responses contributed independently to the measured BOLD signal, without interactions between constituent populations (Fig. 7A). In the nonlinear normalization model, these Gaussian responses were augmented with suppressive surrounds, enabling the positive, in-phase drive associated with one stimulus to be modulated by the negative, anti-phase suppressive drive of the competing stimulus (Fig. 7B). Because this suppressive influence varies with the oscillating competing stimulus, it dynamically modulates the response to the first stimulus, producing nonlinear mixing of the two driving frequencies. We simulated responses to concurrent stimulation at *f_1_* and *f_2_* under both models. The feedforward model reproduced fundamental responses at both frequencies, including multiplexed responses where both stimulus representations contributed to the same voxel, but did not generate IM. In contrast, the normalization model generated an additional IM component at the difference frequency (*f_2_* − *f_1_*), despite this frequency being absent from the physical inputs. Moreover, simulated IM amplitude increased with the degree of sub-voxel overlap between stimulus representations, reaching its greatest magnitude where both populations contributed strongly to the modeled voxel. Thus, whereas independent feedforward summation could reproduce simultaneous encoding of both stimulus frequencies, the generation of IM required nonlinear suppressive interactions, with its amplitude scaling with the degree of suppression between populations tuned to competing stimuli.

These simulations yielded a specific prediction for the empirical BOLD responses: if IM arises from suppressive interactions between competing stimulus representations, its amplitude should increase with the degree to which both representations contribute to a given voxel. Specifically, IM should be strongest in multiplexing vertices, where both stimulus drives are represented, and weaker in vertices dominated by a single stimulus preference. To test this prediction, we quantified the amplitude of BOLD percent signal change at the IM frequency across empirically identified vertex groups. Consistent with the normalization model, IM amplitude was significantly greater in multiplexing vertices than in *f_1_*- or *f_2_*-selective vertices (Wilcoxon rank-sum tests: *P* two-tailed < 0.05 for all pairwise comparisons; Fig. 7C).

Together, these findings establish two complementary results. First, our simulations demonstrate that suppressive interactions between constituent neuronal populations can generate IM responses that are absent under independent feedforward summation. Second, the empirical BOLD responses exhibit the predicted dependence on overlapping stimulus representations, with IM amplitudes greatest where stimulus competition is expected to be strongest. In conjunction with the spatial organization of enhancement and suppression revealed by response phase, these findings support a normalization-like mechanism linking stimulus-specific cortical responses to their nonlinear interactions.

## Discussion

We demonstrate that multiplexed frequency-tagged fMRI (ft-fMRI) enables spatially resolved measurement of concurrent stimulus-specific BOLD responses, identifying cortical populations selectively encoding individual stimuli as well as those multiplexing concurrent inputs. In addition, multiplexed ft-fMRI identifies nonlinear intermodulation (IM) responses at frequencies absent from the stimulus inputs and distinguishes spatially organized populations responding in-versus out-of-phase with the stimulus drive. Frequency-encoded and phase-resolved maps were reproducible within individuals and robust across 3T and 7T MR field strengths. By jointly considering response phase and IM components, we further demonstrate how multiplexed ft-fMRI can constrain computational models of interactions between competing cortical representations, using normalization as a mechanistic test case.

By assigning distinct temporal frequencies to concurrent stimuli, a fundamental advance of our ft-fMRI platform is the ability to distinguish simultaneous stimulus representations within the time series of a single cortical vertex. ft-fMRI thus separates the responses of differently tuned neuronal populations even when their signals are mixed beyond the spatial resolution of fMRI acquisition. This capacity extends the interpretive scope of fMRI beyond mapping aggregate stimulus preferences toward characterizing the composition and interactions of cortical populations to multiple concurrent driving stimuli.

Our findings build on a long history of frequency-tagging in electrophysiology and M/EEG, where stimulus-specific responses and nonlinear IM components have provided powerful measurements of concurrent neural processing ^13–15,17,36^. However, invasive electrophysiology provides limited spatial coverage for mapping cortical organization, while spatial mixing in M/EEG complicates the localization and phase interpretation of signals arising from closely intermingled cortical populations ^18,19^. Frequency-tagging has also been successfully extended to fMRI. Important early work demonstrated the detection of stimulus-driven BOLD responses at frequencies up to 0.75 Hz ^26^, followed by the sensitive and reproducible localization of face-selective cortical responses using fast periodic stimulation ^29^. These latter two studies established the feasibility and sensitivity of recovering periodic responses from fMRI, but did not address the simultaneous representation and interaction of independently tagged inputs. Our multiplexed ft-fMRI platform bridges these approaches by combining spectral separation with the spatial specificity needed to localize distinct stimulus representations and the nonlinear interactions between them.

Response phase adds an important dimension to this framework. Whereas frequency-tagged amplitude captures the strength of stimulus coupling, phase captures the sign of the coupling, distinguishing responses that vary with versus against the corresponding stimulus drive. The observed spatially organized target–surround arrangement of these opposing responses is consistent with stimulus-driven enhancement and suppression, respectively ^39^. Critically, ft-fMRI can resolve these responses independently for concurrent stimuli, revealing an organization that may be obscured by spatial mixing in M/EEG. Phase therefore provides information about the sign and spatial organization of stimulus coupling that is not available from response magnitude alone.

The utility of these complementary measurements depends on their reliability. Fundamental, multiplexed, IM, and phase-resolved responses exhibited reproducible, individual-specific spatial organization, with fundamental and IM oscillations directly observable in BOLD time series from acquisitions as short as three minutes. Their spatial distributions were also consistent across the tested field strengths and frequency pairings, supporting the practicality of ft-fMRI for investigating concurrent cortical dynamics.

The convergence of phase and IM responses offers a new approach to studying cortical normalization. Conventional fMRI investigations have inferred normalization from subadditive responses to concurrent stimuli relative to their separately presented components ^24,25,43^. Although consistent with divisive normalization, such effects are inferred from response differences across experimental conditions rather than from measurements of the interaction itself. Multiplexed frequency-tagging instead isolates nonlinear spectral components during concurrent stimulation, while phase reveals a spatial organization of positive and negative stimulus coupling that may provide the substrate for these interactions. Our sub-voxel pRF model linked these complementary observations mechanistically. Consistent with normalization, we showed that suppressive interactions were required to generate IM responses, and their predicted spatial dependence was supported by the empirical measurements. Thus, multiplexed ft-fMRI lays the groundwork for non-invasively investigating normalization with more complex stimuli, in other sensory modalities, and during cross-modal interactions ^25,44^.

Moving further up the cortical hierarchy, multiplexed ft-fMRI could inform models of attention, decision-making, and memory ^1–7,45^. For example, we have recently deployed this technique to explore how directed attention reshapes stimulus-specific responses, suppressive organization, and nonlinear interactions within the same cortical populations ^41^. Enrichment of multiplexed, intermodulation and phase information into multivariate pattern analyses may reveal how interactions between concurrent stimuli shape distributed population codes ^25,46–48^. Neurodevelopmental and neurodegenerative disorders, ranging from autism ^10^ to Alzheimer’s disease ^49^, are thought to directly impact the circuit basis of normalization. Multiplexed ft-fMRI provides a computationally principled framework for examining normalization, and the downstream consequences of altered competitive processing, in these conditions.

Multiplexed ft-fMRI was also motivated by the need for better translation of findings from computational neuroscience and non-human animals models to non-invasive human neuroimaging. Because individual stimulus drives can be recovered without requiring separate behavioral reports for each input, multiplexed ft-fMRI could facilitate direct comparisons of cortical computations across species ^50–52^ and may be useful in human imaging settings where traditional task fMRI is challenging or infeasible, including neonatal and infant fMRI ^53–56^, and patients with disorders of consciousness ^57,58^.

Several limitations warrant consideration. First, BOLD signals are indirect measures of neuronal activity and are subject to hemodynamic filtering, which reduces sensitivity at higher stimulation frequencies and limits access to rapid synaptic dynamics ^20,26,59^. More systematic characterization of the relationship between frequency spacing, response sensitivity, and hemodynamics will be important for optimizing the ft-fMRI framework. Second, although opposing phase relationships are consistent with stimulus-linked enhancement and suppression, BOLD phase does not directly establish the underlying neuronal mechanism, and regional differences in hemodynamic timing may influence phase estimates. Likewise, IM responses establish nonlinear mixing in the measured response but cannot independently distinguish among neuronal circuit interactions, nonlinear neurovascular contributions, or specific mechanisms such as lateral inhibition and feedback modulation ^14^. The normalization model therefore provides a mechanistic account consistent with the observed responses rather than a unique identification of their circuit origin. Future combinations with electrophysiology, biophysical modeling, and layer-sensitive fMRI could help distinguish the neural and vascular contributions to frequency- and phase-resolved responses.

In summary, our multiplexed ft-fMRI platform combines the temporal separation of concurrent stimulus representations with the spatial specificity needed to resolve their cortical organization. By integrating stimulus-specific responses, phase information, and nonlinear intermodulation components with computational modeling, this framework extends the reach of fMRI from mapping cortical organization to interrogating the neuronal computations that arise from interactions among concurrently active neural populations.

## METHODS

### RESOURCE AVAILABILITY

#### Lead contact

Further information and requests for resources should be directed to and will be fulfilled by the lead contact, [Reebal Rafeh] ().

#### Materials availability

This study did not generate new unique reagents or biological materials.

#### Data and code availability

The code for preprocessing the fMRI data is available at https://github.com/gnngo4/fastfmriprep, and the code for conducting frequency-tagging analyses is available at https://github.com/gnngo4/fastfmri_toolbox. Supplemental movies and information for the simulations can be found at https://osf.io/rqegz/files/osfstorage

### EXPERIMENTAL MODEL AND SUBJECT DETAILS

#### Human participants

10 young adult participants (4 female, self-reported) with normal or corrected-to-normal vision participated as paid volunteers after giving informed written consent. Their ages ranged from 22-29 years old (M = 26.1, SD = 3.49). Participants were compensated at a rate of CAD$15 per hour for their participation in the study. Participants also received an additional CAD$0.05 for each correct response during the behavioral task to incentivize their completion of all experimental sessions. All experimental procedures were approved by the Health Sciences Research Ethics Board at Western University.

A total of 25 frequency-tagging experiments were conducted across the 3 sets of experiments: 3T, 7T, and 3T/7T Vary. This included 9 experiments at 3T, 4 at 7T, and 12 in the 3T/7T Vary group (6 per field strength across 2 participants). A detailed breakdown is provided in Supplementary Table 1.

All participants in the 3T experiment performed both a ‘frequency-tagging’ and a ‘control condition’. 1 participant from the 3T experiment performed the Vary experiment.

## METHOD DETAILS

### Experimental design

The initial goal of this study was to validate a dual frequency paradigm by tagging two frequency-encoded populations using fast fMRI. To achieve this, the first set of experiments, labeled as 3T experiments, included two task conditions: frequency-tagging and control. In the frequency-tagging condition, the stimulus modulated the intensity of two visual-field quadrants using two different frequencies, referred to as fundamental frequencies (see Fig. 1a to see the stimulus schematic). In contrast, the control condition uses the same set-up with frequencies set to 12 Hz and could not be sampled by fMRI. Upon analyzing the dataset, we found evidence for other BOLD dynamics that appeared to be a product of nonlinear mixing of the fundamental frequencies referred to as intermodulation. Since intermodulation frequencies are a function of the fundamental frequencies, a subset of experiments, labelled as 3T/7T Vary experiments were conducted where one of the two stimulated frequencies were modulated across three task conditions and in two subjects. Each subject had a different fundamental frequency modulated. These additional experiments were designed to assess whether the intermodulation frequency-encoded maps were in fact modulated by the underlying fundamental frequencies. Additional experiments were also conducted using a 7T MRI to assess the generalizability of these results at a higher MR field strength. A total of 25 frequency-tagging experiments were conducted across the 3 sets of experiments: 3T, 7T, and 3T/7T Vary. This included 9 experiments at 3T, 4 at 7T, and 12 in the 3T/7T Vary group (6 per field strength across 2 subjects). A detailed breakdown is provided in Supplementary Table 1.

Sample size was determined based on prior fMRI studies mapping responses to single frequency-tagged visual stimuli e.g. ^26,59^ and was selected to provide sufficient within-participant data to reliably characterize frequency-specific responses. No formal a priori power analysis was performed. Because experimental conditions and frequency-tagged responses were defined objectively by the stimulus and analysis procedures, data collection and analyses were not performed blind to the experimental condition.

For analyses involving frequency-responsive cortical vertices, vertices were included if they exhibited a significant response (*P_unadjusted_* < 0.05) in at least 80% of bootstrap samples (fractional overlap > 0.8). Among vertices meeting this criterion, responses were further classified according to the sign of their response phase as either in-phase, reflecting stimulus-driven activation, or anti-phase, reflecting stimulus-driven deactivation. These vertex-selection and response-classification criteria were applied prior to subsequent statistical analyses.

### Dual frequency stimuli

We generated the experimental stimuli and tasks using PsychoPy3^60^. Stimuli were projected at 60 Hz onto a screen (resolution of 1024×768 pixels) fixed at the back of the scanner bore from a frame rate synchronized stimulus laptop. Participants viewed the projected visual stimuli via a mirror fitted into the head coil placed in front of the eyes (total distance to screen is 84 cm).

We presented participants with a pair of radial wedge stimuli occupying one visual field quadrant and another pair of radial wedge stimuli occupying equal spatial portions of the opposing visual field quadrant^13^. Each wedge was part of a concentric grating pattern. The wedge stimuli extended from 1.5° to 10° eccentricity with a spatial frequency of 2.5 cycles per degree visual angle. Participants initiated each experimental run via a button press. During a 14s baseline period at the beginning of each experimental run, we used red lines to cue our participants to covertly attend to a pair of wedges in one visual field quadrant. Following cue offset, participants were required to detect transient color changes lasting for 500 ms in the cued visual field quadrant. Either one of the attended static radial wedge stimuli changed color to a hue of red at 2-5 s pseudo-random intervals drawn from a uniform distribution. The participant had 750 ms to respond to the color change occurrence via button press with the right index finger. If the response did not occur within the response interval, the response was recorded as a false alarm. If the participant did not respond at all before the next color change appeared, this was recorded as a missed trial.

Color change intensity was adjusted for each participant to achieve a response accuracy of approximately 80%, where response accuracy was calculated as the number of correct responses divided by the total sum of correct responses, false alarms and missed trials.

The radial wedge stimuli presented in the unattended visual quadrant were sine-squared modulated at 0.125 Hz and 0.2 Hz (frequency-tagging condition) or did not oscillate (control condition).During both conditions, the unattended stimuli exhibited a 12 Hz counterphase flicker. Each participant from the 3T protocol performed a total of twenty-four 240s runs of each of the frequency-tagging and control conditions over the course of the fMRI sessions, where the stimulus presentation lasted for 219s. For the 7T protocol, participants performed a total of 16 functional runs of the frequency-tagging condition, For the Vary experiment, at each of 3T and 7T, participants performed a total of 12 functional runs for each frequency combination. We counterbalanced the location of the stimuli in either the upper right or upper left visual hemifield across participants. We further counterbalanced the position of the 0.125 Hz and 0.2 Hz stimuli in the upper (left or right) quadrant across participants.

### Choice of stimulation frequencies

A central design question when employing ft-fMRI is what are the best frequencies to stimulate at? Theoretically, the ideal range of stimulation frequencies (to avoid influences of aperiodic and physiological artefacts while retaining sensitivity) lies in the range of 0.1-0.2 Hz, which correspond to the higher end of the aperiodic (1/f) signal range in fMRI (<0.1Hz)^61^, and the lower end of the respiratory range (0.25-0.4Hz)^20,26^, respectively. These stimulus-independent sources of signal variability are apparent when inspecting the run-wise power spectra in Fig. 7. Furthermore, using a stimulation frequency higher than the respiratory range comes at the cost of extreme reduction in sensitivity to frequency-tagged signals^26^. Thus, we intentionally selected stimulation frequency pairs in the range of 0.1-0.2 Hz while ensuring that the stimulus frequency pairs are not (sub)harmonics of one another (e.g. 0.1 Hz and 0.2 Hz), and that we are able to disentangle the frequency-specific signals given the frequency resolution of our experiment (0.006 Hz).

### MRI protocol

All MR imaging was conducted at the Center for Functional and Metabolic Mapping at Western University, using either a 3T or 7T scanner.

### 3T Protocol

Participants in the 3T experiments were scanned on a whole-body 3T MRI scanner (MAGNETOM Prisma, Siemens Healthineers, Erlangen, Germany) equipped with a 64-channel head/neck RF coil. Anatomical imaging was performed only during the first session using a 1 mm isotropic MPRAGE sequence with the following parameters: repetition time (TR) of 2300 ms, inversion time (TI) of 900 ms, echo time (TE) of 2.98 ms, and a flip angle of 9°. GRAPPA acceleration (factor of 2) was applied in the phase-encoding direction, with a total scan duration of 5 minutes. In subsequent sessions, only functional BOLD imaging was conducted. This consisted of 18 runs using a multi-band single-echo gradient-recalled echo (GRE) sequence. Coronal slices were oriented over the occipital pole to specifically target the calcarine sulcus, capturing a limited brain region. Each run lasted 240 seconds and captured 800 volumes. Imaging parameters included: 16 slices at 2.5 mm isotropic resolution, multi-band factor of 4, TR of 300 ms, TE of 30 ms, flip angle of 20°, bandwidth of 2480 Hz/pixel, echo spacing of 0.61 ms, and a right-left phase-encoding direction. Between the 9th and 10th functional runs, a whole-brain field-of-view (FOV) EPI sequence and a GRE field mapping sequence were acquired. While the field map was not used in the analyses, the whole-brain EPI sequence captured 80 slices at 2.0 mm isotropic resolution. This sequence facilitated inter-session registration by aligning small FOV functional images to the anatomical reference via intermediate registration with the larger FOV EPI images.

### 7T Protocol

Participants in the 7T experiments were scanned on a head-only 7T MRI scanner (MAGNETOM 7T MRI Plus, Siemens Healthineers, Erlangen, Germany) equipped with an AC84 Mark II head-gradient coil. Functional imaging was performed using an 8-channel transmit (Tx), 32-channel receive (Rx) RF coil optimized for occipital-parietal (OP) imaging. This coil minimized visual obstruction and increased the signal-to-noise ratio (SNR) in the visual cortex by approximately 35% compared to a whole-head coil.

Anatomical imaging, however, used a whole-head 8-channel Tx, 32-channel Rx RF coil. For anatomical reference, a single 0.7 mm isotropic MP2RAGE scan was acquired during the first session with the following parameters: TR of 6000 ms, first inversion time of 800 ms, second inversion time of 2700 ms, TE of 2.74 ms, flip angles of 4° and 5° for the two inversion pulses, GRAPPA acceleration factor of 3 in the phase-encoding direction, and a slice partial Fourier of 6/8. The scan duration was 10 minutes and 12 seconds. The functional imaging protocol for the 7T experiments was identical to that used in the 3T experiments, consisting of 18 runs of multi-band single-echo GRE BOLD imaging, along with a whole-brain EPI sequence and a GRE field mapping protocol.

### MRI data processing

MRI data was preprocessed using a pipeline developed with *Nipype*, which used the following software packages: *ANTS*, *Freesurfer, FSL*, *AFNI*, *workbench*, and customized *Nipype* workflows adapted from *smriprep* and *fmriprep*. Existing *Nipype* workflows from *fmriprep* were customized to integrate session-level whole-brain EPIs and run-level, narrow FOV EPIs into the registration workflow.

To preprocess each fMRI dataset, slice-timing correction, followed by single-shot interpolation to T1w space, surface resampling to 32k vertex fsLR surfaces (fsLR 32k surfaces), and nuisance regression were performed. fMRI data was regressed using 24 motion parameters (six HMC parameters and their 1^st^ and 2^nd^ order first derivatives[1]), high pass filter regressors (<.01 Hz), and the mean WM and CSF signal to minimize the effects of scanner drift, motion and other non-neural physiological noises. Each fMRI run was truncated between 39 and 219 seconds (or 14 seconds after the oscillatory stimulus begins and when the stimulus ends) to account for transient hemodynamic effects associated with the onset of a continuous oscillatory stimulation.

Details pertaining to resampling of individual fMRI runs to fsLR 32k surfaces are as follows: The subject-specific T1-weighted (T1w) structural image was skull-stripped using *SynthStrip* and then processed with *smriprep* to segment the grey matter (WM), white matter (GM), and cerebrospinal fluid (CSF). Subsequently, the T1w image was registered to the single-band reference (SBRef) image from each scan-specific whole-brain EPI using a boundary-based registration (BBR) cost function. The cortical ribbon mask was transformed into the whole-brain EPI space for each session and utilized to perform a BBR-based registration between the SBRef images of the scan-specific whole-brain EPIs and their corresponding scan- and-run-specific slab EPIs. N4 bias-field corrections were performed to each image prior to performing the registrations. Next, after applying low-pass filtering to each fMRI dataset, head motion correction (HMC) was performed using FSL’s MCFLIRT, registering all volumes to their associated run-specific SBRef image. We found that low-pass filtering removed unwanted fluctuations (0.2-0.4 Hz) in the estimated HMC parameters of a majority of fMRI runs. Note that low-pass filtered data was used solely to generate fluctuation-free HMC transformations, which were then applied to the raw, unfiltered fMRI data. To enable one-shot interpolation of each fMRI run to T1w space, the transformations were concatenated in the following order: HMC to run-specific slab EPI, slab EPI to session-specific whole brain EPI, and whole brain EPI to subject-specific T1w image. Next, the volumetric BOLD data (in T1w space) was resampled into the subject’s native surface, and subsequently resampled onto a standard fsLR 32k surface using spherical surface-based alignment.

### Fitting a frequency-encoded population

To model a frequency from fMRI data, vertex-wise BOLD time series were analyzed using a general linear model (GLM). The GLM included sine and cosine regressors at the frequency of interest to capture the periodic response. An F-test was applied to jointly assess the contributions of both sine and cosine regressors, testing for the presence of a significant response at the specified frequency. The phase offset relative to the stimulus was computed from the ratio of the coefficient weights of the sine and cosine regressors. A formal description of the GLM is provided below.

For any given BOLD time series, the GLM can be expressed as follows:

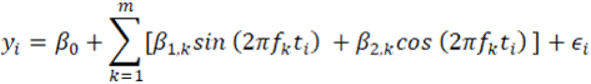

where:

- *^β^*_0_ is the intercept term
- *^β^*_1,*k*_ and *^β^*_2,*k*_ are the coefficients associated to the sine and cosine terms of the k^th^ frequency *^f^_k_*, respectively
- *^f^_k_* is the k^th^ frequency of interest
- ε*_i_* is the error term for observation *^i^*
- *^m^* is the number of frequencies of interest

Additionally, the phase offset (*^v^_k_*) of a frequency of interest can be calculated from the fitted coefficients corresponding to the Fourier basis of that frequency. To align the phase offset with the experimental setup, a constant (π/2) is added to account for the stimulus beginning at the trough (minimum) of the sinusoid. This adjustment ensures that the phase calculation accurately reflects the latency of the BOLD response with respect to the stimulus.

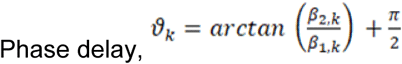

Phase offset was applied as a supplementary filter for identifying frequency-encoded populations. For each frequency of interest, phase delay histograms were constructed across all active vertices and pooled over 3T and 7T experiments using a bin count of 100. Vertices surrounding the histogram peak with a radius of ±^π/2^ were designated as in-phase^29,32^, while all remaining vertices were classified as out-of-phase. In-phase offset ranges were independently estimated for each frequency pair and subject. For example, a separate phase delay histogram was constructed for each frequency pairing experiment across 3T and 7T MR field strengths.

In this study, two sets of frequencies were modelled. First, to assess the feasibility of detecting only fundamental frequency-encoded populations, only *f_1_* and *f_2_* were included in the GLM. Second, to evaluate the presence of intermodulation frequencies, two fundamental (i.e., *f_1_* and *f_2_*) and four intermodulation frequencies (i.e., *f_2_*-*f_1_*, *f_1_*+*f_2_*, 2*f_1_*-*f_2_*, 2f_2_-*f_2_*) were included in the GLM.

### Monte Carlo frequency mapping

#### GLM-based mapping

A Monte Carlo random subsampling approach was employed to generate robust frequency-encoded brain maps. For each participant and task condition (frequency-tagging and control), 400 subsamples of the data were drawn by randomly selecting 50% of preprocessed fMRI runs each time (12/24 runs). The average fMRI signal was computed for each subsample and normalized to %BOLD change.

To generate a frequency-encoded map, a GLM with frequency-specific predictors was applied to each subsample (see Methods section *Fitting a frequency-encoded population* for details). Maps of each k^th^ frequency in the set were binarized by thresholding with an *P_unadjusted_*<.05 (F-test). These activation maps were aggregated across all 400 subsamples by computing the mean, producing a fractional overlap map that indicates the stability of frequency-encoded brain maps across all random subsamples. A value of zero indicates the frequency was not encoded in any of the 400 subsamples, while a value of one indicates it was encoded in all 400 subsamples. For this study, a fractional overlap of 0.8 provided higher sensitivity in areas proximal to regions where the fractional overlap was 1. All activation maps are therefore presented using a fractional overlap of 0.8 with *P_unadjusted_*<.05 to maintain sensitivity to frequency-encoded signals. This choice was especially critical for detecting much weaker intermodulation frequency responses. Vertices exhibiting suprathreshold responses (fractional overlap > 0.8) to only one tagged frequency (*f_1_* or *f_2_*) were classified as frequency-specific vertices, whereas vertices exhibiting suprathreshold responses to both frequencies (*f_1_* and *f_2_*) were classified as multiplexing vertices.

In the 3T/7T frequency pairing experiments, two subjects underwent three frequency-varied conditions across two MR field strengths, with frequency stimulation (*f_1_* and *f_2_*) applied to identical visual fields. Overlap maps were generated for each frequency by creating Monte Carlo random subsampled frequency-encoded maps for each frequency pairing and field strength. The resulting maps were aggregated to create overlap maps across three distinct frequency pairings under each MR field strength condition, and covering both fundamental and intermodulation frequency-encoded populations. To further quantify the overlap between frequency paired conditions and MR field strengths, Dice coefficients were computed for all pairwise comparisons of task conditions and field strengths.

#### Fourier z-score based mapping

To further assess the robustness of the observed effects, we repeated the Monte Carlo random subsampling procedure using a Fourier z-score–based thresholding approach, as implemented in prior fMRI frequency-tagging studies^29,30^. For each vertex, spectral amplitudes at the frequency of interest were converted to Fourier z-scores by normalizing them relative to the local spectral baseline. Specifically, the mean and standard deviation of this spectral baseline were computed from a set of neighboring frequency bins, excluding the frequencies of interest and their harmonics. A frequency “bin” was defined as that lying within a 0.0625 Hz spectral window centered on the frequency of interest, corresponding to 10 adjacent frequency bins (5 on each side) given a frequency resolution of 0.00625 Hz. Fourier z-score was computed as the difference between the Fourier amplitude at the frequency of interest (*^Αf^*) and the mean amplitude at the adjacent resolved frequencies (*^μfn^*), divided by the standard deviation of adjacent frequencies (*^σfn^*).

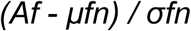

Responses were considered significant if they exhibited an average Fourier Z >3 across subsamples of the data. Vertices exhibiting suprathreshold responses (Fourier Z >3) to only one tagged frequency (*f_1_* or *f_2_*) were classified as frequency-specific vertices, whereas vertices exhibiting suprathreshold responses to both frequencies (*f_1_* and *f_2_*) were classified as multiplexing vertices.

This approach provides a complementary method for detecting frequency-tagged responses by explicitly accounting for the local spectral noise structure. By normalizing the signal of interest by the standard deviation of adjacent frequencies, the Fourier z-score penalizes responses occurring in regions of elevated spectral variance. This is particularly important in fMRI data, where aperiodic low-frequency fluctuations can inflate raw spectral amplitudes. Consequently, Fourier z-score thresholding yields more conservative estimates at lower frequencies, reducing the influence of structured noise and limiting false positives while maintaining sensitivity to true frequency-specific responses.

### Identification of in-phase and anti-phase frequency responses

Suprathreshold (fractional overlap > 0.8) vertex-wise phase offsets were classified as in-phase (stimulus-driven) or out of phase with the stimulus using an iterative procedure to define phase distributions. For each stimulus frequency, the stimulus time course was convolved with a canonical gamma hemodynamic response function (HRF) and truncated to the analyzed time window (39–219 s) to estimate the expected stimulus-driven response phase. We binned response phases and classified the distribution peak with the smallest circular distance to the predicted HRF phase as in-phase. We then used the peak phase value of the distribution as the center phase (φ) and defined all the vertices with their phase values within the window of (φ ± π/2) as part of the in-phase distribution, and vertices with their phase values outside of this window as out of phase with the stimulus ^29^. To ensure robust classification, the procedure was repeated across 500 random subsamples of phase values, and vertices were retained if they were consistently classified as in-phase in ≥95% of resamplings.

Vertices exhibiting significant in-phase synchronization (fractional overlap > 0.8) with the stimulus frequencies were considered feature-tuned. Multiplexing vertices were defined as those showing significant in-phase responses to both stimulus frequencies (*f_1_* and *f_2_*), indicating co-localized responses to simultaneously presented stimuli. Frequency-specific responses were defined as those exhibiting significant in-phase responses to one frequency, but not the other.

### Population receptive field (pRF) mapping

#### pRF mapping experiment

We carried out a population receptive field (pRF) mapping experiment in 3T participants (n=9) during a separate experimental visit. Briefly, the experiment consisted of objects, faces, and words overlaid on pink noise presented through wedge, ring, or bar apertures^62,63^. Participants performed four 300s runs of each of the following three conditions (12 runs total) using the 3T acquisition protocol: 1-‘ccw’ involved a wedge aperture rotating counter-clockwise over the course of the run. 2-‘expand’ involves a ring aperture expanding outward from the center of the visual field. 3-‘multibar’ involves a sliding bar aperture moving horizontally, vertically, and diagonally across the visual field. We averaged the data across runs of the same type (i.e. ccw, expand, and multibar). Stimulus apertures indicating the position of the texture were prepared at a resolution of 768 pixels x 768 pixels. We then used analyzePRF^22^ to triangulate the polar angle and eccentricity of vertex-wise pRFs. Vertices whose pRF model explained more than 10% of the variance (R2 > 0.1) in the time series data during the pRF mapping experiment were selected for further analyses of their visual field coverage.

#### pRF analysis and visual field coverage (VFC)

Visual-field coverage (VFC) plots represent the sensitivity of defined frequency-specific and multiplexing vertices to different locations in the visual field. For each participant, we combined the gaussian receptive field models for each suprathreshold vertex. The summation of these gaussians resulted in a maximum value for each location in the visual field. The resulting visual field coverage plots thus represent the maximum envelope of sensitivity across the visual field for analyzed vertices.

To compute stimulus signed overlap index, we calculated the mean pRF VFC value (defined as the mean value in a specific portion of the visual-field coverage plot) in the visual field regions in which the target frequency-tagged stimuli were presented (overlap_f1_ or overlap_f2_) and computed the difference (overlap_f1_ – overlap_f2_). A positive value thus represents a visual field bias towards the *f_1_* stimulus location, a negative value indicates a visual field bias towards the *f_2_* stimulus location, whereas a visual field bias of 0 indicates co-localization to the *f_1_* and *f_2_* stimulus locations.

#### Variation in IM amplitude with polar distance from diagonal separating frequency-tagged stimuli

We examined spatial variability in IM amplitude by computing the relationship between polar distance from the diagonal between the frequency-tagged *f_1_* and *f_2_* stimuli. We selected suprathreshold IM (*f_2_* - *f_1_*) vertices (fractional overlap > 0.8) with pRFs within 45° of the diagonal for each participant. We obtained the estimated BOLD amplitude from the GLM fit for selected IM vertices, and aggregated IM amplitude values across participants. We binned vertices between −45° and 45° in 12 steps (7.5°s per step) and obtained the mean amplitude in each bin.

#### Significance testing of population overlaps between fundamental and intermodulation frequencies

To account for the spatial autocorrelation inherent in frequency-encoded populations, a generative modeling approach was employed to examine spatial dependencies between any two populations. For each frequency-encoded population, an unthresholded fractional overlap map was used to generate a surrogate fractional overlap map that preserved the spatial autocorrelation structure. This surrogate map was binarized by thresholding to retain only the top *n* vertices, matching the vertex count of the original frequency-encoded population. This process produced a single binarized surrogate frequency-encoded map. To construct null distributions for an overlap metric, this procedure was repeated 500 times, resulting in a set of surrogate maps.

The degree of overlap between two populations was quantified using the Dice coefficient, with alternative measures such as the Jaccard coefficient yielding similar results. Overlap between two spatial maps was deemed significant if the observed overlap score exceeded the 95^th^ percentile of the overlap scores derived from the null surrogate maps. Significance was assessed for each subject and task condition, allowing for comparisons between the topography of fundamental-frequency-encoded populations and intermodulation-encoded populations.

Since multiple fundamental frequency-encoded populations can exhibit significant overlap with an intermodulation population, subsequent analyses focused on determining which fundamental population predominantly composed the IM population. Because IM frequencies arise from the nonlinear combination of *f_1_* and *f_2_*, the neural population encoding them was expected to overlap more strongly with the multiplex population than with individual fundamental frequency population alone. To evaluate this, a one-sided Wilcoxon signed-rank test was conducted to assess whether overlap scores between the IM population and the multiplex population were significantly greater than those between the intermodulation and each fundamental frequency population (*f_1_*, *f_2_*). Statistical analyses of population overlaps and differences were performed across all 25 frequency-tagging experiments.

#### Dynamic models of frequency-tagged population responses

We created a population receptive field (pRF) model which captured the activity of differentially tuned subpopulations within fMRI voxels. Each pRF consisted of either homogeneously tuned subpopulations (i.e. those responding to only one of the two stimulus locations) or heterogeneously tuned subpopulations tuned to either of two frequency-tagged stimulus locations. We constructed classic subpopulation receptive fields (sRFs) using Gaussian distributions to represent the stimulus drive at different spatial locations. The stimulus drive at a given location was determined by the contrast of the stimulus at that location, where contrast was modulated over time as a sine squared function of the stimulus frequency. Thus, at a given timepoint*t*, the classic sRF is given by

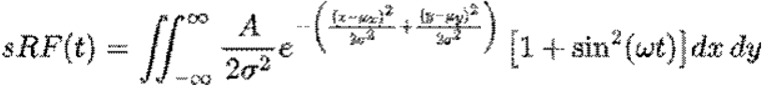

where

- *^A^* is the amplitude of the Gaussian
- ^σ^ is the standard deviation of the Gaussian
- ^μ^*_x_*^,μ^*_y_* is the mean (center point) of the distribution on the *^x^* and *^y^* axis
- ω is the angular frequency calculated as ^2^*_πf_*, where *^f^* ω is the stimulus frequency

In addition, we constructed sRFs using a Difference of Gaussians (DoG) distribution to capture both the stimulus drive and surrounding suppressive drive as per the normalization model, where at a given timepoint, the normalization model sRF is given by

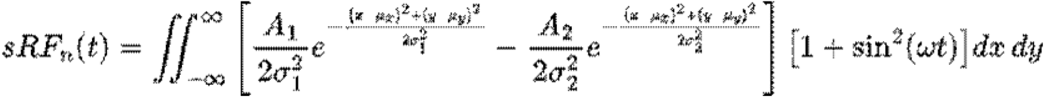

where

- *^A^_1_^,A^_2_* are the amplitudes of the narrow (central) Gaussian and the wider (surround) Gaussian, where *^A^_1_^≥A^_2_*
- *^σ^_1_^,σ^_2_* are the standard deviation of the central and surround Gaussians, where *^σ^_1_^≤σ^_2_*
- ^μ^*_x_*^,μ^*_y_* is the mean (center point) of the distribution on the *^x^* and *^y^* axis
- ^ω^ is the angular frequency calculated as ^2*πf*^, where *^f^* is the stimulus frequency

In turn, the net activity of the pRF at a given timepoint is calculated as the sum of sRF activities encompassed by that pRF, meeting the zero activation threshold, producing a rectified signal

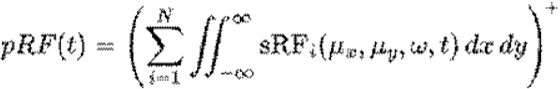

The index *^i^* determines the position and frequency parameters ^μ^*_x_*^,μ^*_y_*^,ω^ of each sRF contained in the pRF.

#### Eye-tracking acquisition & analysis

Eye tracking was performed using an Eyelink 1000 system during the experiment. Eye tracking data was sampled at 0.5 kHz. For each experimental run, we obtained a time series of horizontal and vertical eye positions. Blinks were automatically detected by the Eyelink system and manually selected based on outlier values, and data within ±0.1 seconds of each blink were excised. We detrended recorded eye-positions to account for instrumental drift over the course of the experimental session. We plotted a 2D histogram of horizontal and vertical eye-positions aggregated across experimental runs. Eye positions were summarized by fitting a 2D Gaussian probability distribution to the data and calculating the area of the contour that contained 95% of the fitted distribution (Supplementary Fig. 1)

#### EEG acquisition & analysis

We sampled EEG activity at 512 Hz using a 32-channel BioSemi system and referenced the recorded data to active and passive amplifier channels during acquisition. EEG data was then detrended, notch filtered to remove line noise at 60 Hz, and re-referenced to the average of the mastoid channels using the Fieldtrip toolbox^64^. We manually rejected and removed independent components exhibiting blink or muscle-activity artifacts from the EEG data. We discarded the initial 39s from each run prior to analysis to avoid transient effects of signal onset and align the time series of the EEG data to that acquired using fMRI.

We obtained the counterphase amplitude envelope for the run-average signal of each channel by applying a 4th order Butterworth filter to bandpass the signal between 23.8 Hz and 24.2 Hz and then calculating the magnitude of the Hilbert-transform of this signal. We divided the resulting EEG time series into periods corresponding to that of the presented stimuli (i.e. 8s for the 0.125 Hz stimulus and 5s for the 0.2 Hz stimulus) and averaged the data across periods. We plot the average stimulus-locked response envelope for the central occipital EEG channel (Oz) at each frequency. We averaged the maximum amplitude of the counterphase envelope for each channel across participants and plotted these values onto the standard 32-channel BioSemi electrode coordinates in Fieldtrip. We then examined the latency between measured fMRI and EEG stimulus-locked average activity by computing the cross-correlation between the mean fMRI signal across suprathreshold vertices and the down-sampled EEG signal envelope for each participant at lags between ±5s (for the 0.2 Hz stimulus) and ±8s (for the 0.125 Hz stimulus) (Supplementary Fig. 6).

## QUANTIFICATION AND STATISTICAL ANALYSIS

### Population overlap significance

To account for the spatial autocorrelation inherent in frequency-encoded populations, a generative modeling approach was employed to examine spatial dependencies between any two populations. For each frequency-encoded population, an unthresholded fractional overlap map was used to generate a surrogate fractional overlap map that preserved the spatial autocorrelation structure. This surrogate map was binarized by thresholding to retain only the top n vertices, matching the vertex count of the original frequency-encoded population. This process produced a single binarized surrogate frequency-encoded map. To construct null distributions for an overlap metric, this procedure was repeated 500 times, resulting in a set of surrogate maps. The degree of overlap between two populations was quantified using the Dice coefficient, with alternative measures such as the Jaccard coefficient yielding similar results. Overlap between two spatial maps was deemed significant if the observed overlap score exceeded the 95th percentile of the overlap scores derived from the null surrogate maps. Significance was assessed for each subject and task condition, allowing for comparisons between the topography of fundamental-frequency-encoded populations and intermodulation-encoded populations. Since multiple fundamental frequency-encoded populations can exhibit significant overlap with an intermodulation population, subsequent analyses focused on determining which fundamental population predominantly composed the IM population. Because IM frequencies arise from the nonlinear combination of *f_1_* and *f_2_*, the neural population encoding them was expected to overlap more strongly with the multiplex population than with individual fundamental frequency population alone. To evaluate this, a one-sided Wilcoxon signed-rank test was conducted to assess whether overlap scores between the IM population and the multiplex population were significantly greater than those between the intermodulation and each fundamental frequency population (*f_1_*, *f_2_*). Statistical analyses of population overlaps and differences were performed across all 25 frequency-tagging experiments.

### Intermodulation response amplitude

To examine differences in IM amplitude between multiplexing and frequency-specific vertices, we obtained the run-average BOLD signal for each suprathreshold vertex (i.e. fractional overlap > 0.8). We then computed the Fourier amplitude of the vertex signal using the Fast Fourier Transform function from the scipy library. Finally, we averaged the Fourier amplitude at the IM frequency across vertices for each vertex group.

### Run and session reliability

Frequency-encoded maps of *f_1_*, *f_2_* and *f_2_*-*f_1_* were generated at the run level (*P_unadjusted_*<0.05) to assess the reliability of these maps. For each experiment (3T, 7T, and frequency pairing), intra- and inter-subject Dice overlap scores were computed to evaluate the consistency of maps at the run and session levels across experiments. In the 3T experiment, conditions were further divided into 3T Left and 3T Right, corresponding to stimulation in the left and right hemifields, respectively. This separation ensured that overlap score comparisons were not confounded by task differences. Pairwise Dice overlap scores were calculated independently for each experiment, comparing maps across all subjects at both the run and session levels. To account for varying FOVs associated with different frequency-encoded maps, comprehensive masks were created at both the run and session levels for each experiment. These masks identified vertices common to all FOVs, enabling consistent estimation of overlap scores. Overlap scores were categorized as intra-subject (overlap between maps from the same subject) or inter-subject (overlap between maps from different subjects). To determine whether frequency-encoded maps were subject-specific, statistical tests were applied. The Kolmogorov-Smirnov (KS) test compared the distributions of intra- and inter-subject Dice scores, while the Mann-Whitney U (MW-U) test compared their means. To visualize frequency-tagged dynamics, run-level maps for a single subject were aggregated by calculating the overlap across runs. A fractional overlap of 0 indicated vertices with no overlap across runs, while a value of 24 indicated vertices present across all runs. A threshold of 0.5 was applied to create a region of interest (ROI) for extracting time series data from both run-level and session-averaged fMRI data for that subject. Vertex-level time series data were converted to %BOLD, followed by Z-score normalization and displayed as vertically stacked carpet plots, showing session-level and run-level dynamics for the *f_1_*, *f_2_* and *f_2_*-*f_1_* frequencies. To enhance visualization of frequency-locked bands specific to each frequency population, contributions from the other two frequencies were regressed out using a general linear model. For example, visualizing *f_1_* time series involved regressing out the effects of *f_2_* and *f_2_*-*f_1_*. This approach emphasized the frequency-tagged dynamics of lower-power signals, such as *f_2_* and *f_2_*-*f_1_*. Carpet plots were used to visualize the spectral profiles of both session- and run-level vertices. Power spectra were estimated via Welch’s method, using a segment length of 601 samples with 50% overlap, resulting in a frequency resolution of ∼0.006 Hz. To enhance visibility of peaks at fundamental and intermodulation frequencies, vertex-level power spectra were Z-score normalized.

### Phase map reliability

For each participant, we acquired data for each of our task conditions over the course of multiple experimental sessions. To test for the reliability of the reported phase at each frequency, we inspected the consistency of phase offset across experimental sessions. We first generated session specific phase and maps estimated from the run-average data of each experimental session. These maps were thresholded according to the criteria described in Activation thresholding (i.e. fractional overlap > 0.8). We then examined the circular correlation between thresholded session-specific maps. To test for the significance of phase map correlations across experimental sessions, we employed a generative modeling procedure since surface maps of task metrics are spatially autocorrelated. For each participant, we used each session’s thresholded metric maps to generate 50 simulated maps that retain the spatial autocorrelation structure of the original map. The correspondence between each session’s map with these surrogate maps was then computed to generate a distribution of surrogate circular correlations. We tested for the reliability of subject phase maps by comparing the distribution of between-session map circular correlation values to the surrogate distribution. The reported test statistic was obtained by aggregating the within-participant between-session distributions of map correspondence values and the corresponding surrogate distributions across participants before performing the Kolmogorov-Smirnov (K-S) test to test for differences between these aggregated distributions, and the Mann-Whitney (M-W) U test to test for statistical differences between the distribution means. To examine whether the observed spatial topographies of our phase maps are specific to each participant, we compared within-participant between-session map correlations to between-participant session map correlations. To generate the between-participant distribution, we computed the circular correlation between within-participant session maps and session maps for participants who were presented with the same stimulus configuration (i.e. those presented with the identical positioning of oscillating wedge stimuli in the same visual field quadrant), thresholded using the same mask (fractional overlap > 0.8) applied to the within-participant session maps. We then compared the distribution of within-participant session map correlation values against the distribution of between-participant session map correlation values. The reported test statistics were obtained by aggregating the within-participant distributions and between-participant distributions and performing the K-S test for differences between these distributions, and performing the M-W U test to test for statistical differences between the distribution means.

### Control and frequency-tagging condition comparisons

In the 3T experiment, which involved 9 subjects who completed both a control condition and a frequency-tagging condition, analyses were conducted to evaluate differences in sensitivity between fundamental and intermodulation frequencies. Statistical significance for all analyses was assessed using a one-sided Wilcoxon signed-rank test.

### Statistical analysis

Statistical analysis was performed using the scipy and statsmodels libraries in python. For vertex-wise inference, the statsmodels library was used to fit the GLM to the time series data and perform the F-test to determine the significance of frequency-specific responses. For participant-wise statistical inference, no assumptions were made about the normality of compared data distributions. The scipy library was used to perform the reported non-parametric statistical tests. An alpha level of *P_unadjusted_* < 0.05 was used to assess significance. No multiple-comparisons correction was applied.

## ADDITIONAL RESOURCES

Supplemental movies and simulation information: https://osf.io/rqegz

