## Supplementary Figures for "A frequency-tagged fMRI platform for accessing spatiotemporally resolved cortical computations"

Supplementary Table 1: Summary of Frequency Tagging Experiments

| Experiment | Conditions | Frequencies, Hz | Subjects (IDs) | Sessions/Subject | Runs/Session | Total Runs |
|---|---|---|---|---|---|---|
| 3T | Control & Frequency-tagging | $f_1$=0.125, $f_2$=0.2 | 9 (s000-s009) | 3 | 9 | 27 per condition |
| 7T | Frequency-tagging | $f_1$=0.125, $f_2$=0.2 | 4 (s010-s013) | 3 | 9 | 27 |
| 3T/7T Vary | Frequency-tagging ($f_1$/$f_2$ pairings) | s014:<br>A: $f_1$=0.125, $f_2$=0.2<br>B: $f_1$=0.125, $f_2$=0.175<br>C: $f_1$=0.125, $f_2$=0.15<br><br>s015:<br>D: $f_1$=0.125, $f_2$=0.2<br>E: $f_1$=0.15, $f_2$=0.2<br>F: $f_1$=0.175, $f_2$=0.2 | 2 (s014, s015) | 2 (at both 3T & 7T) | 6 | 12 per condition |

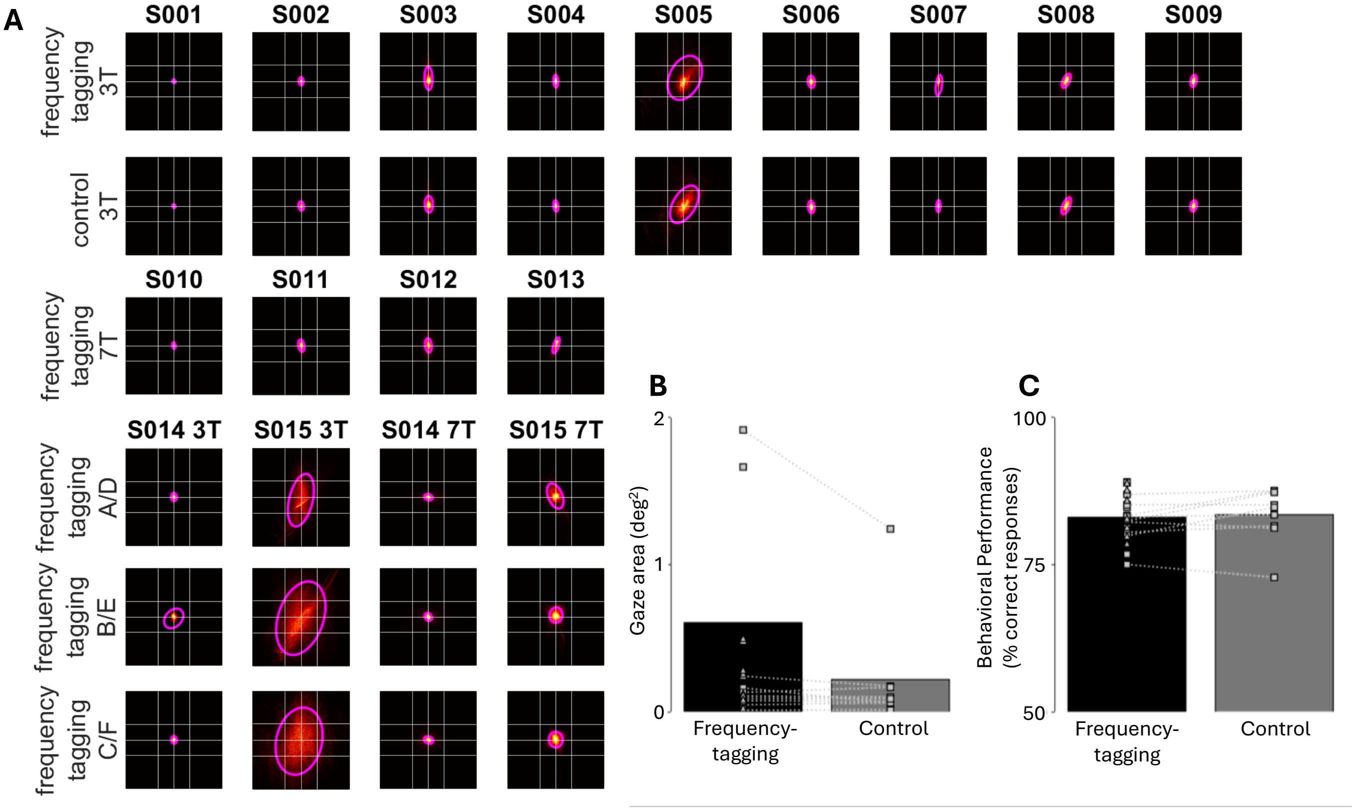


**Supplementary Figure 1. Participant behavioral performance across experimental conditions.**

(A) Eye positions remain consistently near the center of the display. Eye-tracking data is summarized by displaying the heatmaps and fitted gaussian distribution of the aggregated data for each participant (columns) and experimental condition (rows). The grid lines indicate the radius of the central fixation region. The magenta contour lines indicate the area of the gaussian mixture model that contains 95% of the fitted data. Eye-tracker could not be properly calibrated for S004 and S015 at 3T, leading to large deviations in measured gaze positions.

(B) The gaze contour areas are comparable between experimental conditions across participants (KS test; $P$=0.71). Data points indicate the participant mean gaze contour area across runs for an experimental condition. Data points are generally constrained by the area of the central fixation region (grid lines).

(C) Participant behavioral performance was comparable between experimental conditions across participants (KS test; $P$=0.46). Data points indicate the participant mean behavioral performance across runs for an experimental condition. These data show that behavioral performance was maintained at ~80%.

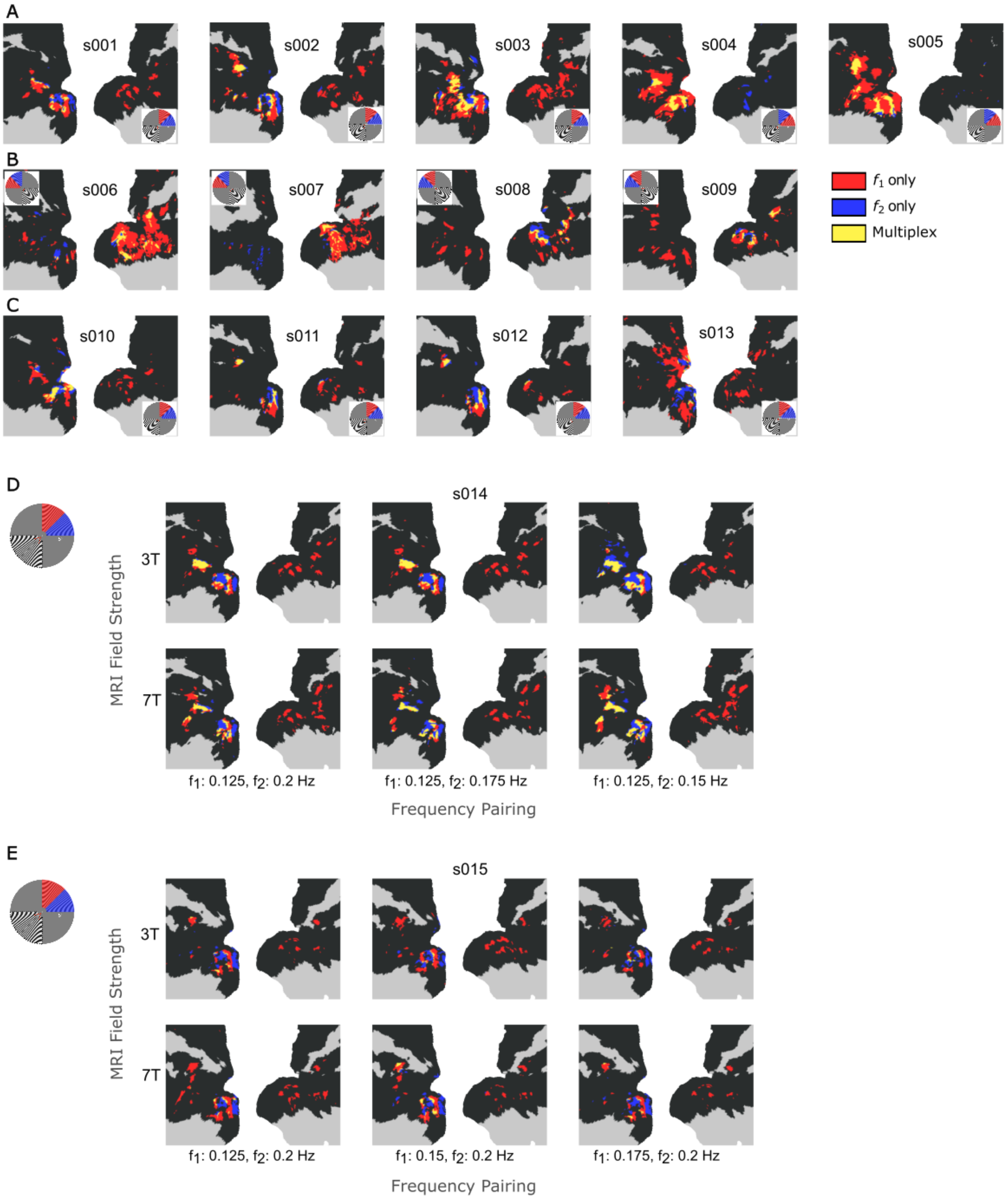


**Supplementary Figure 2. Individual subject fundamental frequency-encoded maps (GLM approach)**

Frequency-encoded maps were estimated in individual participants using a Monte Carlo subsampling procedure applied to the frequency-tagging condition (*n* = 400 subsamples). Vertices were considered significant at $P < 0.05$ and retained if identified in ≥ 80% of subsamples. Red and blue indicate vertices encoding $f_1$ and $f_2$, respectively, and yellow indicates vertices exhibiting multiplexed responses to both frequencies. Insets show the stimulated visual field locations for $f_1$ (red) and $f_2$ (blue).

(A) 3T participants with visual frequency-tagged stimuli presented in the upper right visual field quadrant
(B) 3T participants with visual frequency-tagged stimuli presented in the upper left visual field quadrant
(C) 7T participants with visual frequency-tagged stimuli presented in the upper right visual field quadrant
(D) Participant with visual frequency-tagged stimuli presented in the upper right visual field quadrant across different frequency-pairings (columns) and field strengths
(E) Maps follow the same conventions as panel (D), except for a different participant

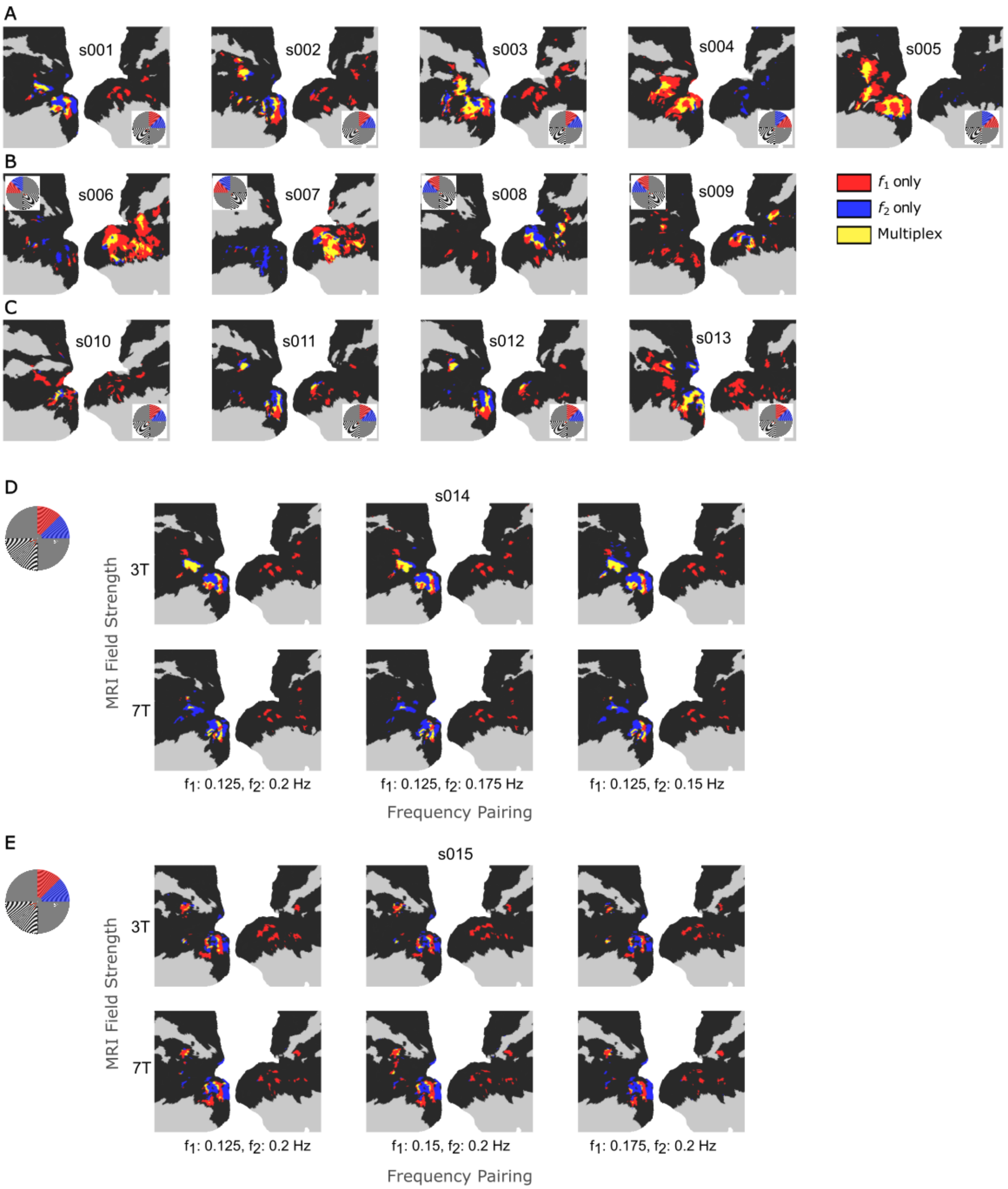


**Supplementary Figure 3. Individual subject fundamental frequency-encoded maps (Fourier z-score approach)**
Frequency-encoded maps were estimated in individual participants using a Monte Carlo subsampling procedure applied to the frequency-tagging condition ($n$ = 400 subsamples). Vertices were considered significant if they exhibited an average Fourier z-score > 3 across subsamples. Red and blue indicate vertices encoding $f_1$ and $f_2$, respectively, and yellow indicates vertices exhibiting multiplexed responses to both frequencies. Insets show the stimulated visual field locations for $f_1$ (red) and $f_2$ (blue).

(A) 3T participants with visual frequency-tagged stimuli presented in the upper right visual field quadrant
(B) 3T participants with visual frequency-tagged stimuli presented in the upper left visual field quadrant
(C) 7T participants with visual frequency-tagged stimuli presented in the upper right visual field quadrant
(D) Participant with visual frequency-tagged stimuli presented in the upper right visual field quadrant across different frequency-pairings (columns) and field strengths
(E) Maps follow the same conventions as panel (D), except for a different participant

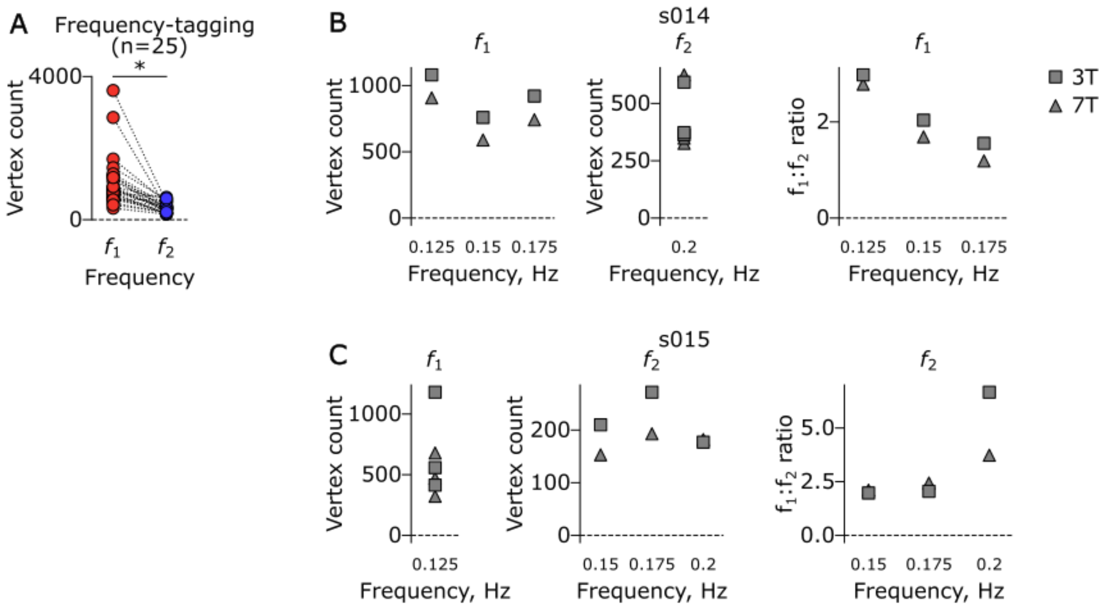


**Supplementary Figure 4. Relationship between stimulation frequency and sensitivity.**

(A) Intra-experiment sensitivity of frequency-encoded populations, across all frequency-tagging experiments where the frequency $f_1<f_2$, showed higher sensitivity in $f_1$ populations.

(B) And (C) Inter-experiment sensitivity of frequency-encoded populations in two subjects (s014 and s015) with varying frequencies ($f_1$ and $f_2$, respectively). In both subjects (left and middle columns for s014 and s015, respectively), a consistent trend of increasing vertex count with decreasing stimulation frequency was not observed across the varied frequencies. To account for inter-experimental effects that may influence the expected trends, the $f_1$:$f_2$ ratio was computed and shown in the right column. A higher ratio indicates that the visual system is more effectively frequency-tagged at a lower frequency ($f_1$), supporting the expectation of higher sensitivity at lower stimulation frequencies.

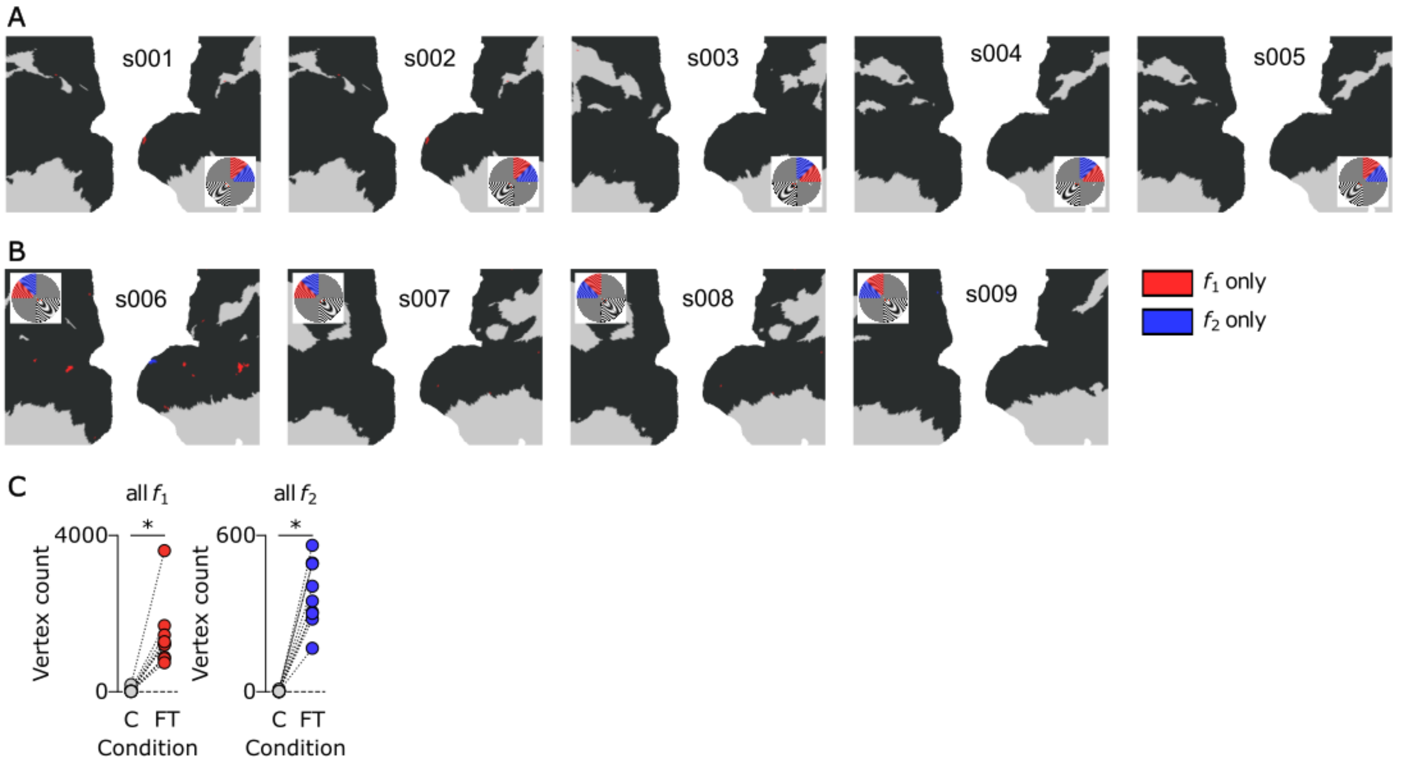


**Supplementary Figure 5. Fundamental frequencies are not detected under a control condition.**

(A) Frequency-encoded maps under a control condition across the 3T experiment. The vertices were identified using a Monte Carlo random subsampling approach. The significance threshold was set at unadjusted *P*<.05 with vertices appearing in at least 80% (320 out of 400) of the random subsamples. In these maps, red denotes the $f_1$ and blue denotes $f_2$. No vertices showing multiplexing (interaction of f1 and f2 vertices) were identified across all 9 subjects in the 3T experiment. Subject-matched maps under the frequency-tagging condition can be seen in main figure 1A-B.

(B) Increased sensitivity to $f_1$ and $f_2$ vertices under the frequency-tagging [or FT] condition, compared to the control [or C] condition. **P*<.05 Wilcoxon signed-rank test

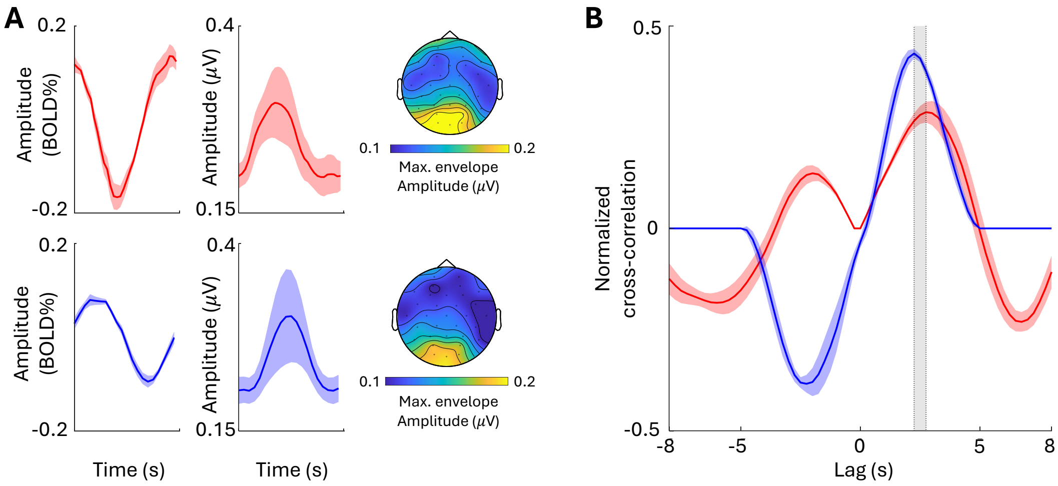


**Supplementary Figure 6. Cross-correlation between stimulus-locked average EEG and fMRI BOLD oscillation (n=8).**

(A) Average stimulus-locked BOLD (left) and EEG (right) oscillations at f1 (top) and f2 (bottom). Shaded region indicates the SE across participants. EEG topographic maps show that the amplitude of stimulus-locked oscillations is maximal around occipital electrodes.

(B) Cross-correlation between EEG and fMRI stimulus-locked oscillations. The phase lag between EEG and fMRI BOLD oscillations fell within a physiologically plausible range of ~2.5 seconds at both stimulation frequencies (shaded gray area).

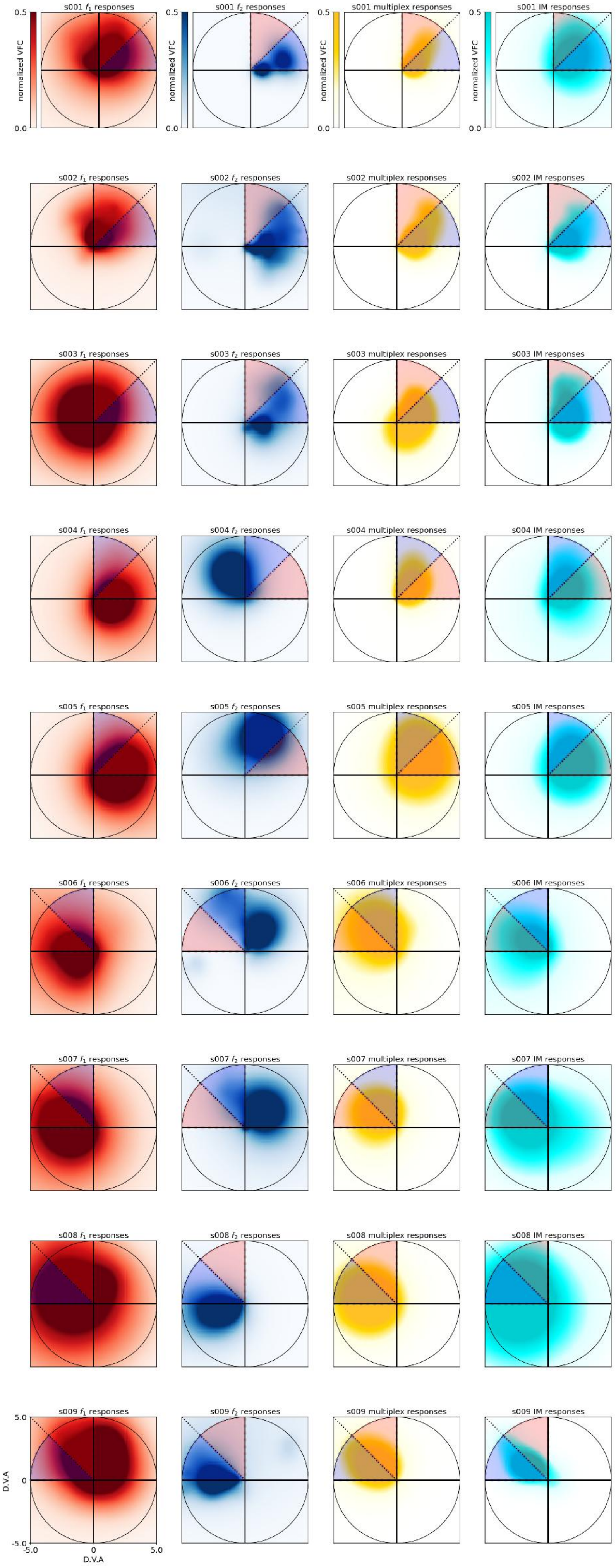


**Supplementary Figure 7. Visual field coverage (VFC) maps of frequency-specific, multiplex, and IM responses for individual participants**

Visual field coverage maps of frequency-specific ($f_1$: red; $f_2$: blue), multiplexing (yellow), and non-linear IM (cyan) responses (columns) for individual participants (rows). Red wedges indicate the position and extent of the $f_1$ checkerboard wedge stimulus for each participant, and blue wedges indicate the position of the $f_2$ wedge for each participant. Normalized visual field coverage (VFC) represents the proportion of pRFs responding to a visual field location. D.V.A is degrees visual angle.

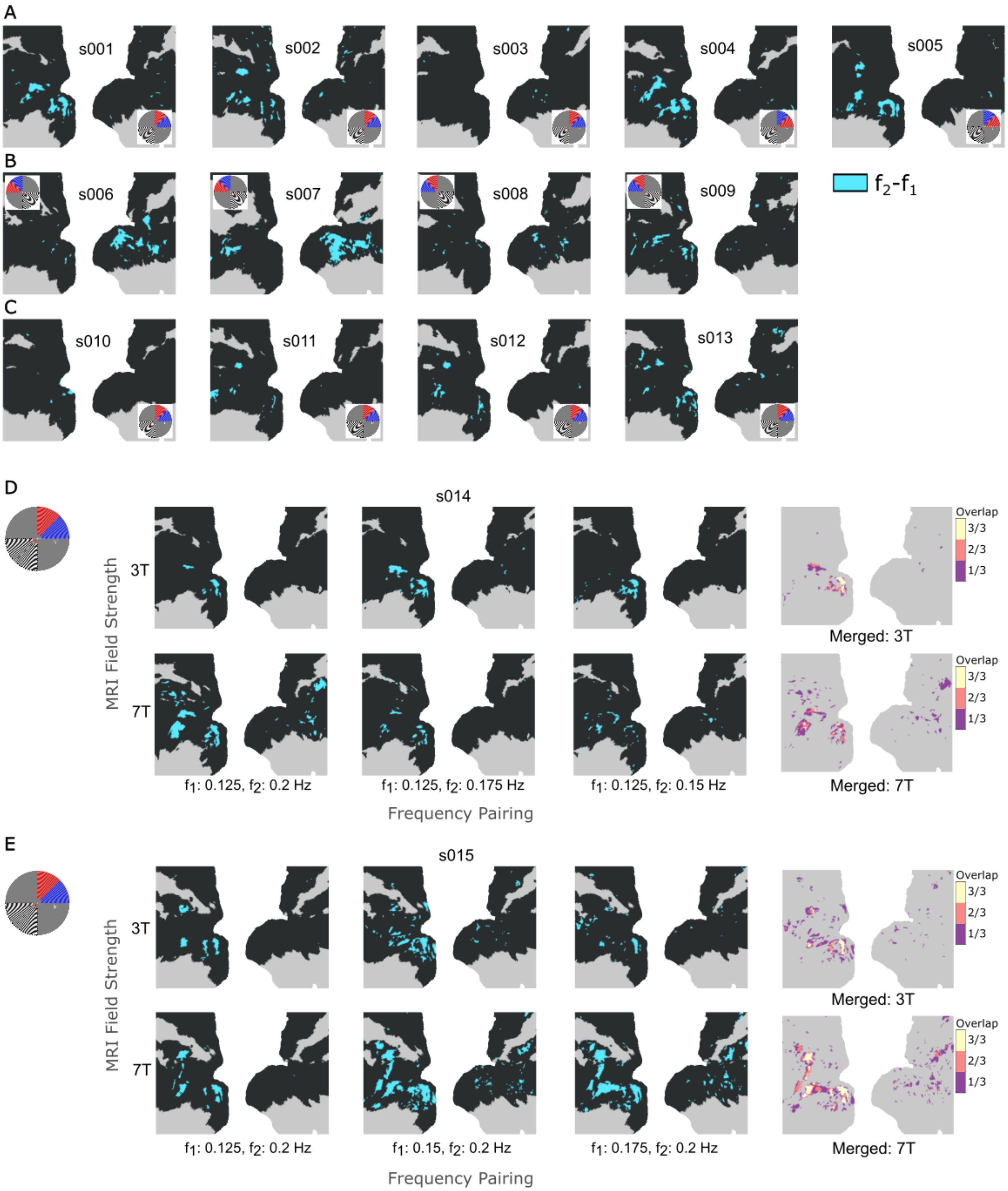


**Supplementary Figure 8. Individual subject non-linear intermodulation frequency-encoded maps (GLM approach)**
Frequency-encoded maps were estimated in individual participants using a Monte Carlo subsampling procedure applied to the frequency-tagging condition ($n$ = 400 subsamples). Vertices were considered significant at $P < 0.05$ and retained if identified in ≥ 80% of subsamples. Cyan indicates vertices encoding $f_2$- $f_1$. Insets show the stimulated visual field locations for $f_1$ (red) and $f_2$ (blue).

A) 3T participants with visual frequency-tagged stimuli presented in the upper right visual field quadrant
B) 3T participants with visual frequency-tagged stimuli presented in the upper left visual field quadrant
C) 7T participants with visual frequency-tagged stimuli presented in the upper right visual field quadrant
D) Participant with visual frequency-tagged stimuli presented in the upper right visual field quadrant across different frequency-pairings (columns) and field strengths
E) Maps follow the same conventions as panel (D), except for a different participant

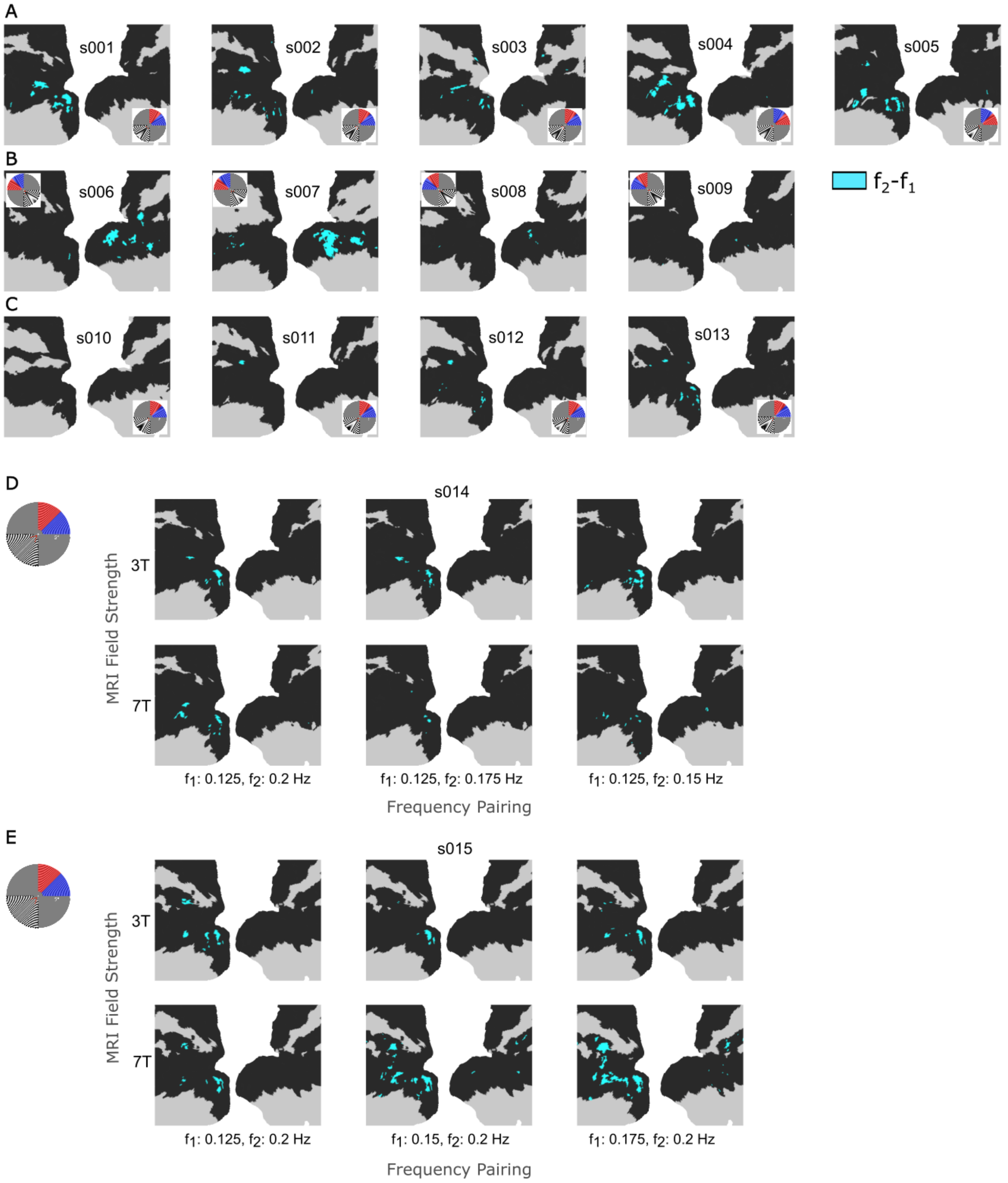


**Supplementary Figure 9. Individual subject non-linear intermodulation frequency-encoded maps (Fourier z-score approach)**

Frequency-encoded maps were estimated in individual participants using a Monte Carlo subsampling procedure applied to the frequency-tagging condition ($n$ = 400 subsamples). Vertices were considered significant if they exhibited an average Fourier z-score > 3 across subsamples. Cyan indicates vertices encoding $f_2$- $f_1$. Insets show the stimulated visual field locations for $f_1$ (red) and $f_2$ (blue).

A) 3T participants with visual frequency-tagged stimuli presented in the upper right visual field quadrant
B) 3T participants with visual frequency-tagged stimuli presented in the upper left visual field quadrant
C) 7T participants with visual frequency-tagged stimuli presented in the upper right visual field quadrant
D) Participant with visual frequency-tagged stimuli presented in the upper right visual field quadrant across different frequency-pairings (columns) and field strengths
E) Maps follow the same conventions as panel (D), except for a different participant

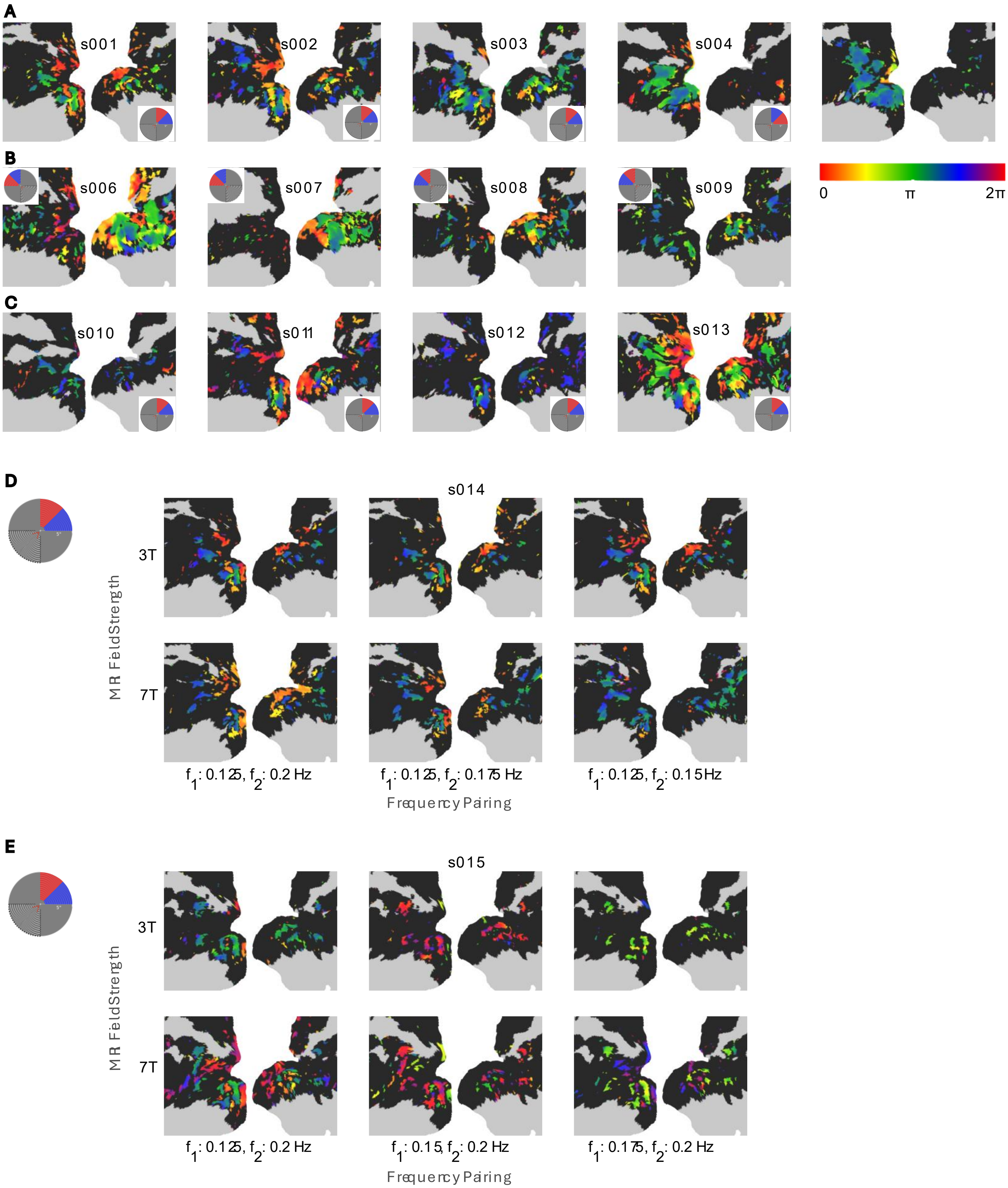


**Supplementary Figure 10. Individual subject *$f_1$* response phase maps**

*$f_1$* frequency-specific phase maps were estimated in individual participants using a Monte Carlo subsampling procedure applied to the frequency-tagging condition (*n* = 400 subsamples). Vertices were considered significant at $P < 0.05$ and retained if identified in ≥ 80% of subsamples. The color bar denotes the response phase scaled to the period of the examined frequency. Insets show the stimulated visual field locations for *$f_1$* (red) and *$f_2$* (blue).

A) 3T participants with visual frequency-tagged stimuli presented in the upper right visual field quadrant
B) 3T participants with visual frequency-tagged stimuli presented in the upper left visual field quadrant
C) 7T participants with visual frequency-tagged stimuli presented in the upper right visual field quadrant
D) Participant with visual frequency-tagged stimuli presented in the upper right visual field quadrant across different frequency-pairings (columns) and field strengths
E) Maps follow the same conventions as panel (D), except for a different participant

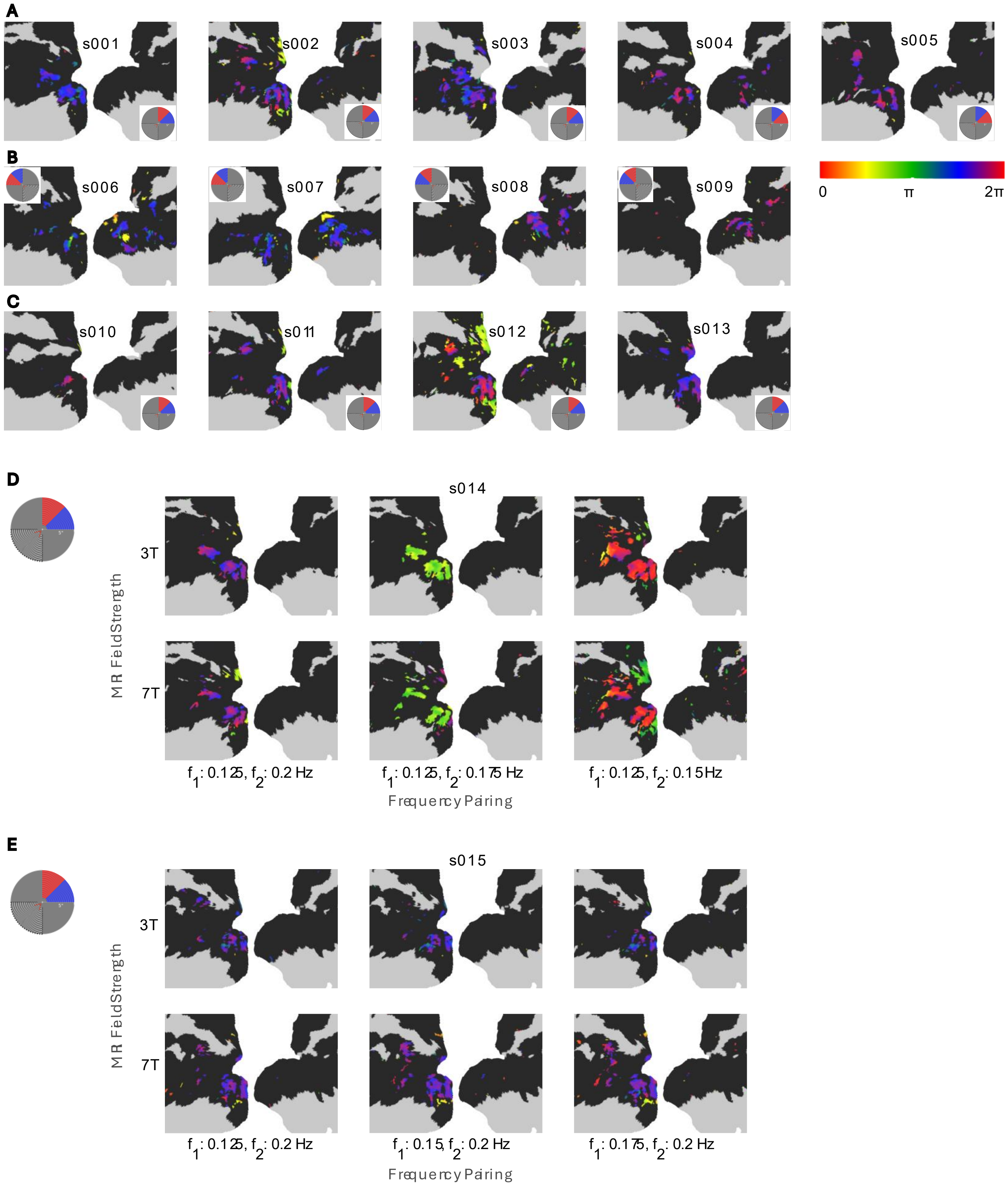


**Supplementary Figure 11. Individual subject *f2* response phase maps**

$f_2$ frequency-specific phase maps were estimated in individual participants using a Monte Carlo subsampling procedure applied to the frequency-tagging condition ($n$ = 400 subsamples). Vertices were considered significant at $P < 0.05$ and retained if identified in ≥ 80% of subsamples. The color bar denotes the response phase scaled to the period of the examined frequency. Insets show the stimulated visual field locations for $f_1$ (red) and $f_2$ (blue).

A) 3T participants with visual frequency-tagged stimuli presented in the upper right visual field quadrant

B) 3T participants with visual frequency-tagged stimuli presented in the upper left visual field quadrant

C) 7T participants with visual frequency-tagged stimuli presented in the upper right visual field quadrant

D) Participant with visual frequency-tagged stimuli presented in the upper right visual field quadrant across different frequency-pairings (columns) and field strengths

E) Maps follow the same conventions as panel (D), except for a different participant

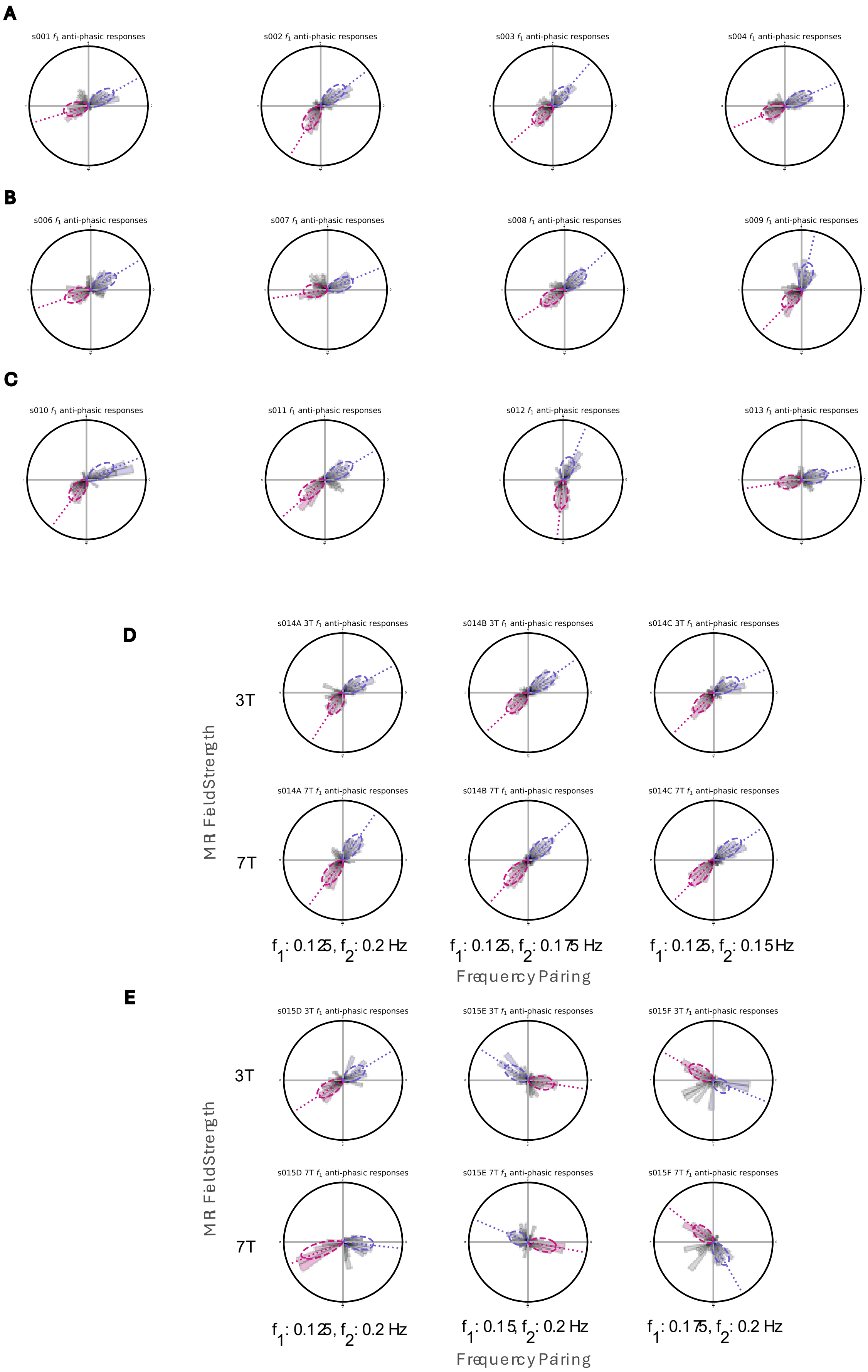


in-phase
anti-phase

**Supplementary Figure 12. Individual subject $f_1$ phase clustered distributions** Frequency-specific suprathreshold response phases were classified into in-phase (violet-red) and anti-phase (slate-blue) synchronized populations based on a resampling procedure. Polar plots display the response phase distribution across suprathreshold cortical vertices. Colored distributions consistently responded in-phase (violet-red) or anti-phase (slate-blue) with the stimulus oscillations across subsamples of the data (>0.95 of subsamples). Gray distributions did not meet the cut-off for consistent samplings. Dotted lines indicate the circular median of the in-phase (violet-red) and anti-phase (slate-blue) distributions. Dashed area indicates the von Mises circular fit to the in-phase (violet-red) and anti-phase (slate-blue) distributions.

A) 3T participants with visual frequency-tagged stimuli presented in the upper right visual field quadrant
B) 3T participants with visual frequency-tagged stimuli presented in the upper left visual field quadrant
C) 7T participants with visual frequency-tagged stimuli presented in the upper right visual field quadrant
D) Participant with visual frequency-tagged stimuli presented in the upper right visual field quadrant across different frequency-pairings (columns) and field strengths
E) Maps follow the same conventions as panel (D), except for a different participant

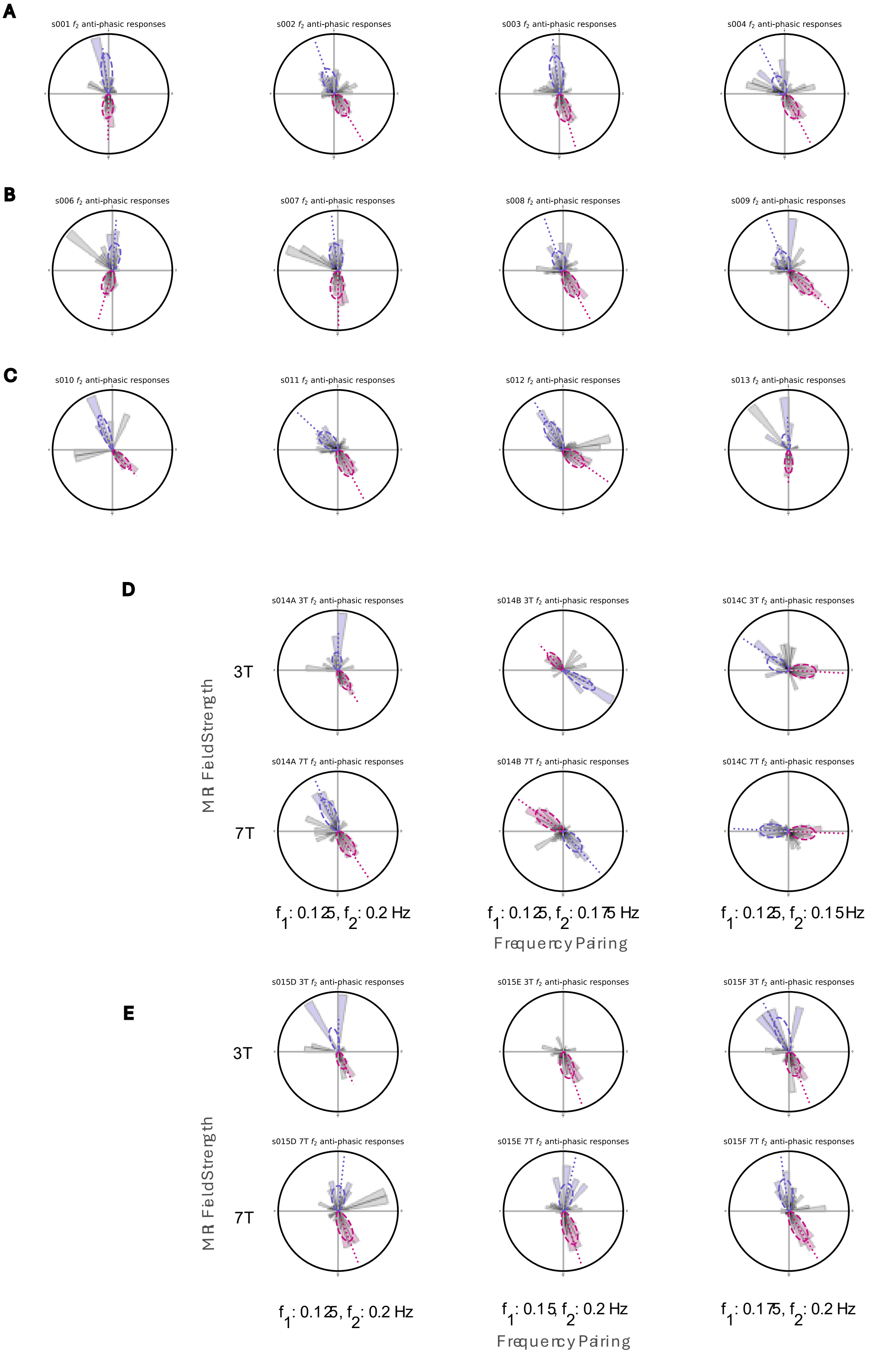


**Supplementary Figure 13. Individual subject $f_2$ phase clustered distributions** Frequency-specific suprathreshold response phases were classified into in-phase (violet-red) and anti-phase (slate-blue) synchronized populations based on a resampling procedure. Polar plots display the response phase distribution across suprathreshold cortical vertices. Colored distributions consistently responded in-phase (violet-red) or anti-phase (slate-blue) with the stimulus oscillations across subsamples of the data (>0.95 of subsamples). Gray distributions did not meet the cut-off for consistent samplings. Dotted lines indicate the circular median of the in-phase (violet-red) and anti-phase (slate-blue) distributions. Dashed area indicates the von Mises circular fit to the in-phase (violet-red) and anti-phase (slate-blue) distributions.

A) 3T participants with visual frequency-tagged stimuli presented in the upper right visual field quadrant
B) 3T participants with visual frequency-tagged stimuli presented in the upper left visual field quadrant
C) 7T participants with visual frequency-tagged stimuli presented in the upper right visual field quadrant
D) Participant with visual frequency-tagged stimuli presented in the upper right visual field quadrant across different frequency-pairings (columns) and field strengths
E) Maps follow the same conventions as panel (D), except for a different participant

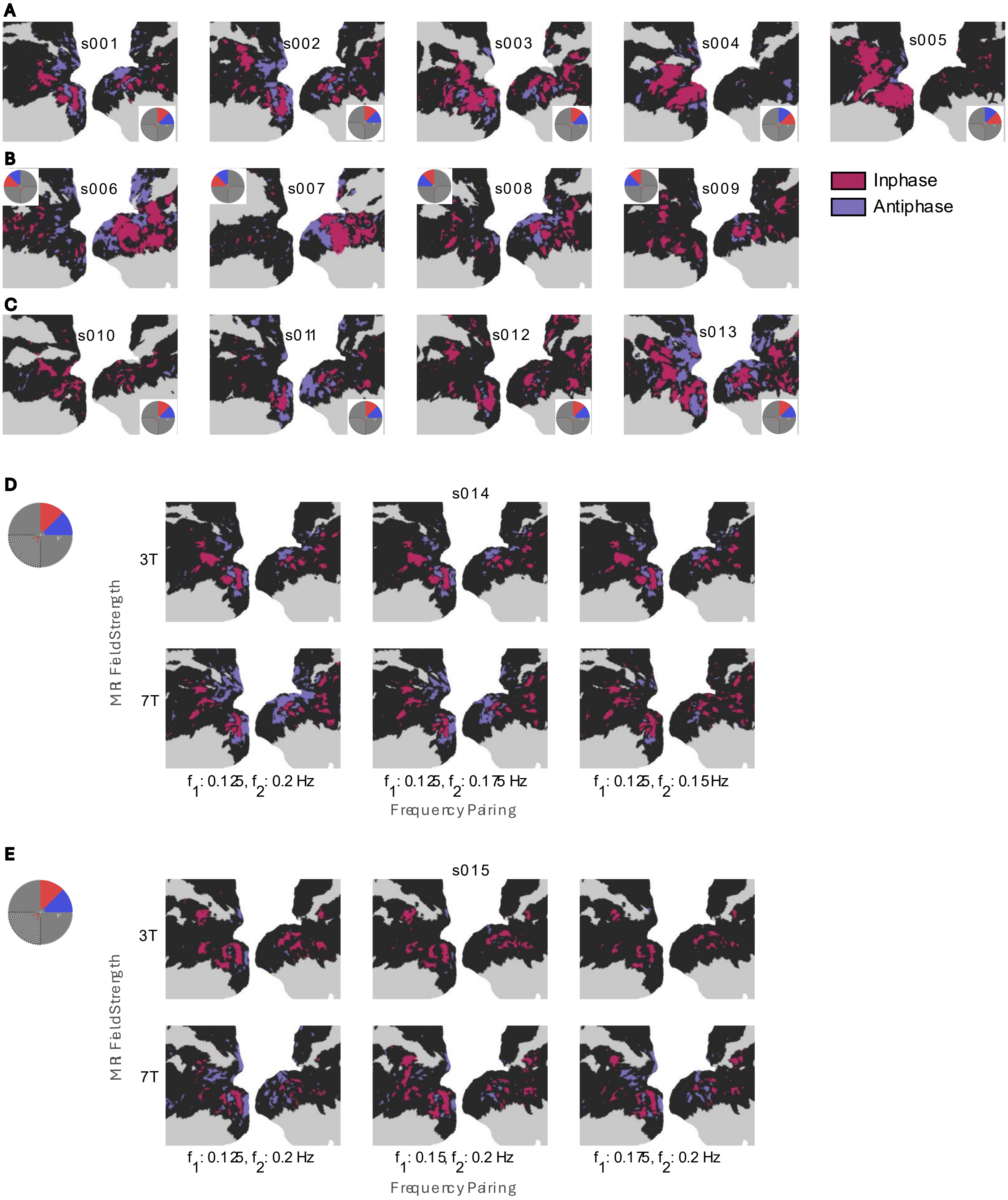


**Supplementary Figure 14. Individual subject $f_1$ signed phase maps**

$f_1$ frequency-specific phase maps were classified into in-phase (violet-red) and anti-phase (slate-blue) synchronized populations based on a resampling procedure. Colored vertices consistently responded in-phase (violet-red) or anti-phase (slate-blue) with the stimulus oscillations across subsamples of the data (>0.95 of subsamples). Insets show the stimulated visual field locations for $f_1$ (red) and $f_2$ (blue).

A) 3T participants with visual frequency-tagged stimuli presented in the upper right visual field quadrant
B) 3T participants with visual frequency-tagged stimuli presented in the upper left visual field quadrant
C) 7T participants with visual frequency-tagged stimuli presented in the upper right visual field quadrant
D) Participant with visual frequency-tagged stimuli presented in the upper right visual field quadrant across different frequency-pairings (columns) and field strengths
E) Maps follow the same conventions as panel (D), except for a different participant

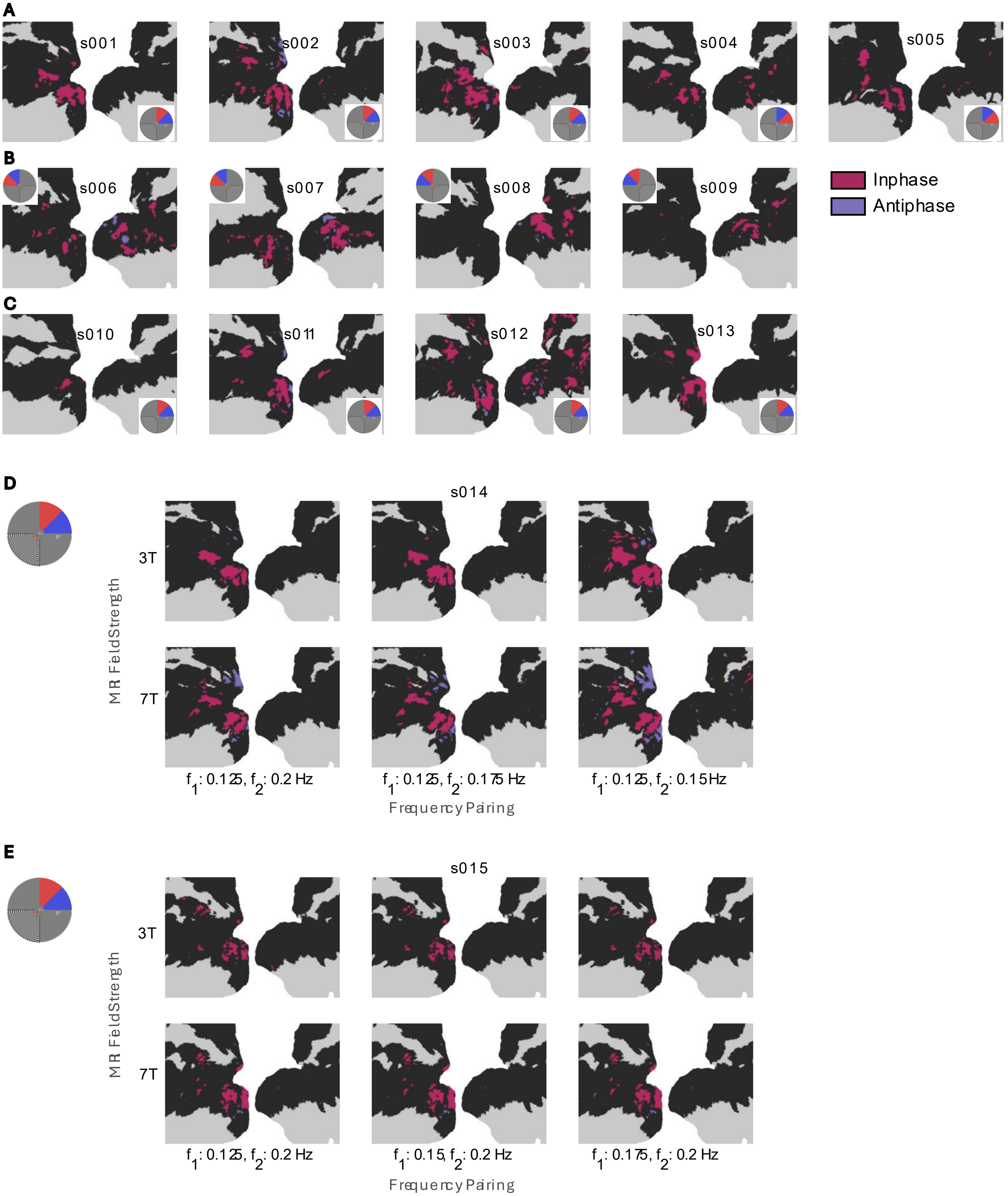


**Supplementary Figure 15. Individual subject $f_2$ signed phase maps**

$f_2$ frequency-specific phase maps were classified into in-phase (violet-red) and anti-phase (slate-blue) synchronized populations based on a resampling procedure. Colored vertices consistently responded in-phase (violet-red) or anti-phase (slate-blue) with the stimulus oscillations across subsamples of the data (>0.95 of subsamples). Insets show the stimulated visual field locations for $f_1$ (red) and $f_2$ (blue).

A) 3T participants with visual frequency-tagged stimuli presented in the upper right visual field quadrant
B) 3T participants with visual frequency-tagged stimuli presented in the upper left visual field quadrant
C) 7T participants with visual frequency-tagged stimuli presented in the upper right visual field quadrant
D) Participant with visual frequency-tagged stimuli presented in the upper right visual field quadrant across different frequency-pairings (columns) and field strengths
E) Maps follow the same conventions as panel (D), except for a different participant

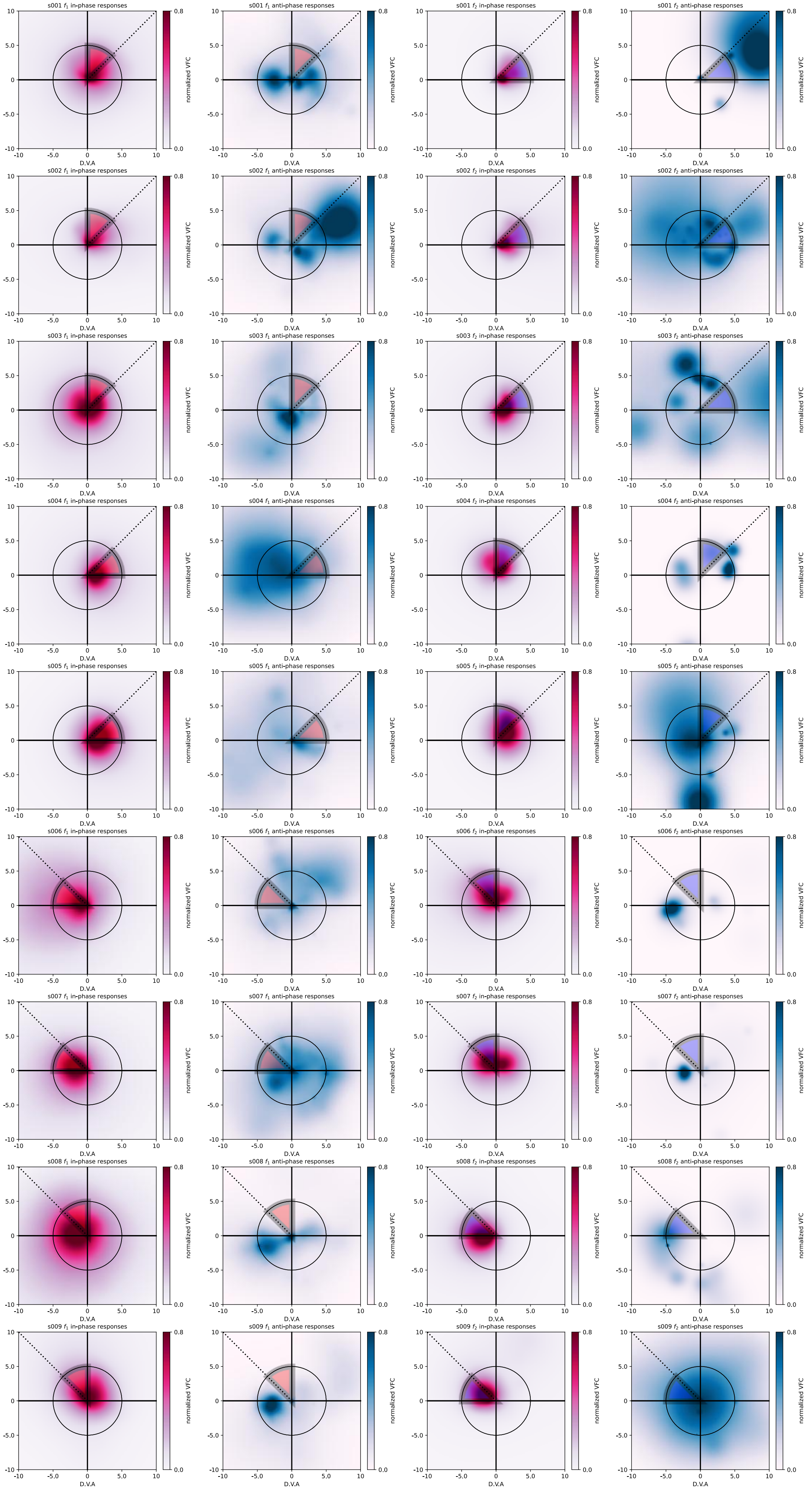


**Supplementary Figure 16. Visual field coverage maps for in-phase and anti-phase responses across participants and frequencies**

Visual field coverage (VFC) maps of suprathreshold in-phase (violet-red) and anti-phase (slate blue) responses across participants (rows; n=9) at each frequency of interest ($f_1$: first 2 columns) and ($f_2$: last 2 columns).

Outlined red wedges indicate the position and extent of the $f_1$ checkerboard wedge stimulus for each participant, and outlined blue wedges indicate the position of the $f_2$ wedge for each participant. Normalized visual field coverage (VFC) indicates the proportion of pRFs responding to a visual field location. D.V.A is degrees visual angle.

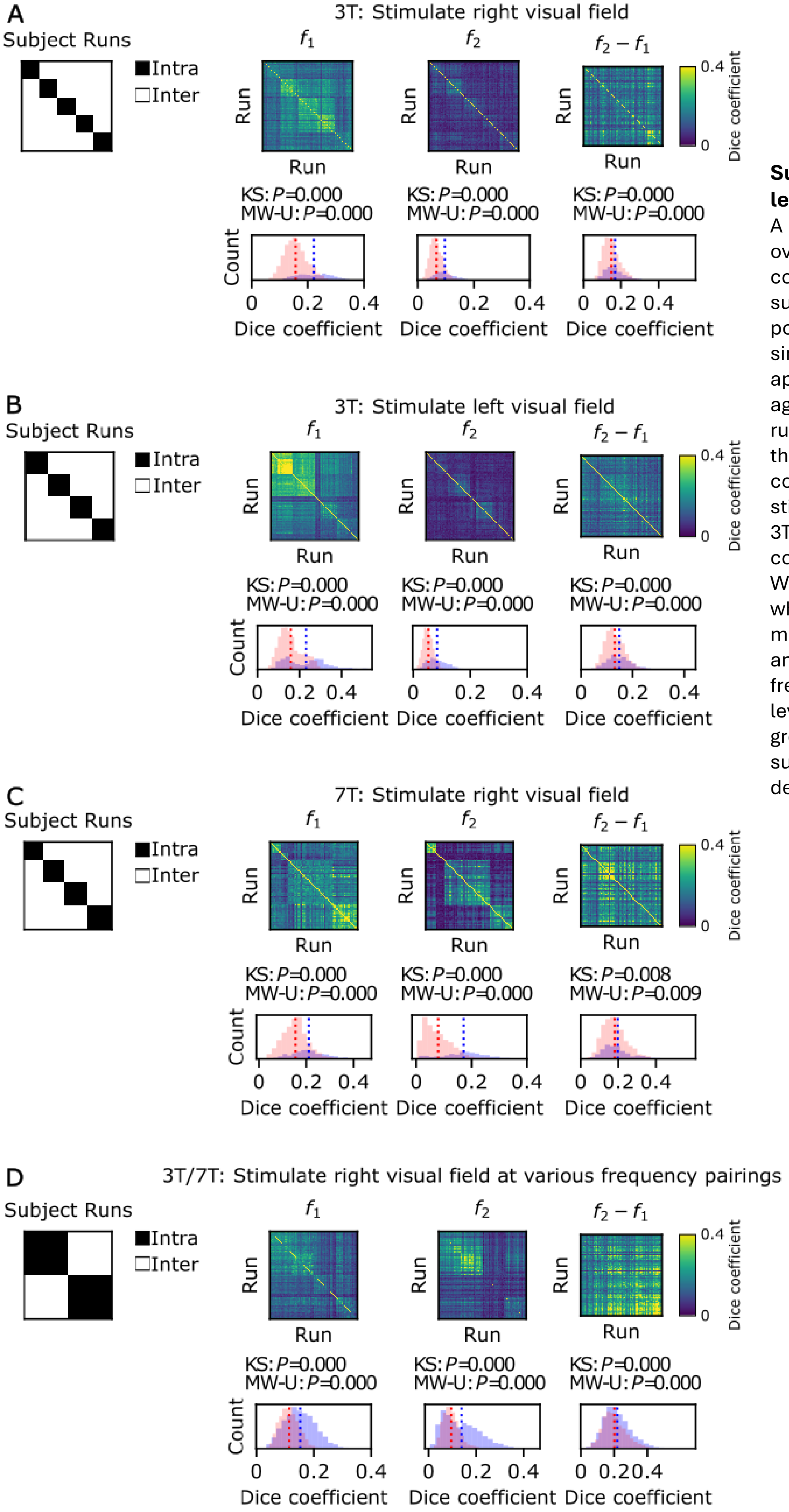


**Supplementary Figure 17. Specificity of run-level frequency-encoded maps.**
A subject's run-level map demonstrates higher overlap with their own run (intra-subject) compared to runs from other subjects (inter-subject) across $f_1$, $f_2$, and $f_2$-$f_1$ frequency populations. Run-level maps were derived from single frequency-tagging fMRI run (no averaging applied). These run-level maps were compared against the subject's other run maps and against run maps from other subjects. (A, B, C, D) show the Dice coefficient across all run-by-run comparisons, controlling for visual-field stimulation location: (A) 3T (stimulate-right), (B) 3T (stimulate-left), (C) 7T, and (D) 3T/7T mixed conditions. Kolmogorov-Smirnov (KS) and Mann-Whitney U (MW-U) tests were used to evaluate whether intra-subject and inter-subject run-level map overlaps significantly differed in distribution and mean, respectively. Results indicate all frequency-encoded populations ($f_1$, $f_2$, $f_2$-$f_1$) run-level maps exhibit subject-specificity, with greater overlap observed within the same subject. *P*-values of KS and MW-U tests are denoted above each plot.

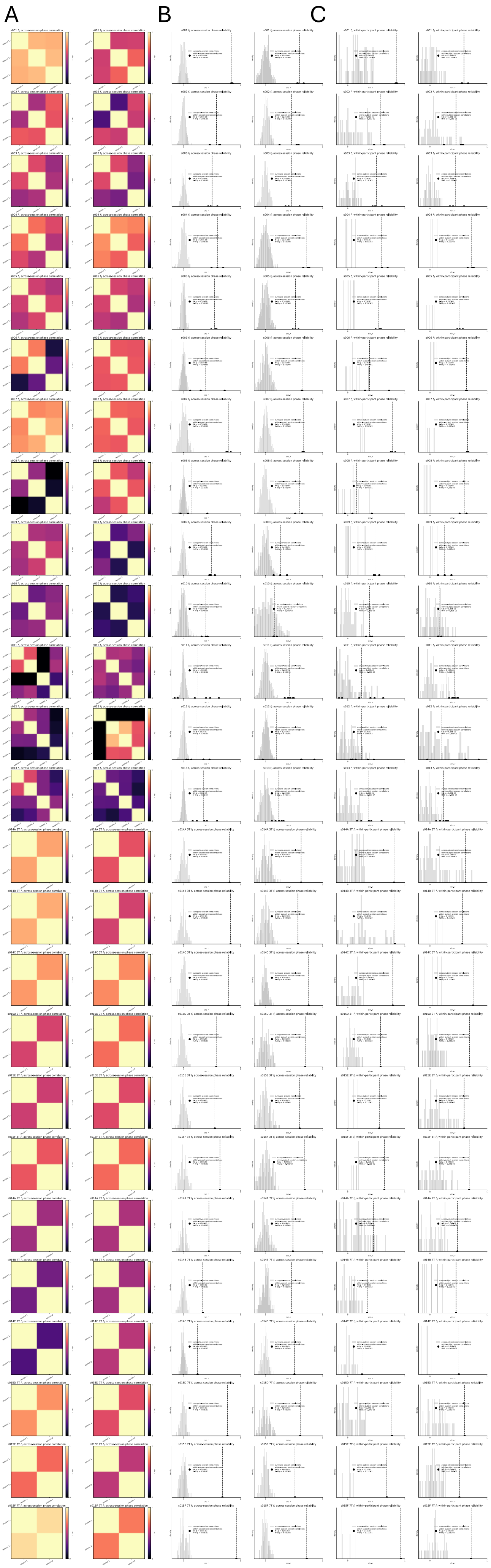


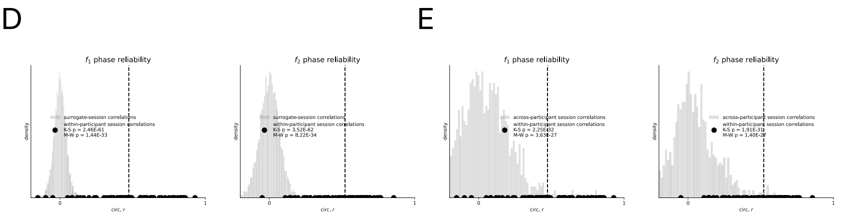


**Supplementary Figure 18: Reliability of phase maps across sessions and within participants**

a) Correlation (circular *r*) between phase maps estimated from each experimental session for each participant (n=15, rows) at $f_1$ (left) and $f_2$ (right).

b) Surrogate map test to determine the significance of between session correlations. Data in gray shows the distribution of surrogate correlation values generated by correlating session phase maps with shuffled phase maps that retained the same spatial autocorrelation structure and anatomical distribution as the original $f_1$ (left) and $f_2$ (right) session phase maps. Black data points represent the between session phase map correlations within participants. Dashed vertical lines indicate the mean value of between session phase map correlations. Reliability of phase maps was tested by comparing the between session correlations against the surrogate distribution for each participant using one-tailed Kolmogorov-Smirnov and Mann-Whitney U tests (p-values for individual participants reported in plots).

c) Between participant session phase map test to examine the uniqueness of the functional topography of within-participant session phase maps. Data in gray shows the distribution of between-participant session phase map correlation values. Data representation follows the same convention as in (b). Individuality of phase maps was tested by comparing the within-participant between session correlations against the between-participant between-session correlations using one-tailed Kolmogorov-Smirnov and Mann-Whitney U tests (p-values for individual participants reported in plots).

d) The distributions of the between session correlations and surrogate distributions aggregated across all participants (Kolmogorov-Smirnov $p_{f1\ \text{one-tailed}} = 2.46\text{x}10^{-61}$; $p_{f2\ \text{one-tailed}} = 3.52\text{x}10^{-62}$; Mann-Whitney U $p_{f1\ \text{one-tailed}} = 1.44\text{x}10^{-33}$; $p_{f2\ \text{one-tailed}} = 8.22\text{x}10^{-34}$).

e) The distributions of the within-participant between session correlations and between-participant between session correlations aggregated across all participants (Kolmogorov-Smirnov $p_{f1\ \text{one-tailed}} = 1.03\text{x}10^{-20}$; $p_{f2\ \text{one-tailed}} = 1.91\text{x}10^{-31}$; Mann-Whitney U $p_{f1\ \text{one-tailed}} = 3.63\text{x}10^{-27}$; $p_{f2\ \text{one-tailed}} = 1.40\text{x}10^{-27}$).
